# Mechanisms of Premotor-Motor Cortex Interactions during Goal Directed Behavior

**DOI:** 10.1101/2023.01.20.524944

**Authors:** Mansour Alyahyay, Gabriel Kalweit, Maria Kalweit, Golan Karvat, Julian Ammer, Artur Schneider, Ahmed Adzemovic, Andreas Vlachos, Joschka Boedecker, Ilka Diester

**Author notes:** These authors contributed equally to this work (Mansour Alyahyay, Gabriel Kalweit).

## Abstract

Deciphering the neural code underlying goal-directed behavior is a long-term mission in neuroscience^1,2^. Neurons exhibiting preparation and movement-related activity are intermingled in the premotor and motor cortices^3,4^, thus concealing the neural code of planned movements. We employed a combination of electrophysiology, pathway-specific optogenetics, phototagging, and inverse reinforcement learning (RL) to elucidate the role of defined neuronal subpopulations in the rat rostral and caudal forelimb areas (RFA and CFA), which correspond to the premotor and motor cortical areas. The inverse RL enabled the functional dissection of spatially intermingled neuronal subpopulations, complementing our pathway-specific optogenetic manipulations and unveiling differential functions of the preparation and movement subpopulations projecting from RFA to CFA. Our results show that the projecting preparation subpopulation suppresses movements, whereas the projecting movement subpopulation promotes actions. We found the influence of RFA on CFA to be adaptable, with the projection either inhibiting or exciting neurons in the superficial and deep CFA layers, depending on context and task phase. These complex interactions between RFA and CFA likely involve the differential recruitment of inhibitory interneurons in the CFA, which is supported by our electron microscopy analysis of the connectivity between these regions. We provide here unprecedented mechanistic insights into how the premotor and primary motor cortices are functionally and structurally interlinked with the potential to advance neuroprosthetics.

**Graphical abstract:** This study provides mechanistic insights into the interactions between the rostral forelimb area (RFA) and the caudal forelimb area (CFA). Specifically, we provide evidence for a differential impact of RFA on CFA depending on the task phase and the targeted CFA layers. RFA contains at least two spatially intermingled subpopulations - one related to movement preparation and one to movement execution. Both subpopulations project to CFA. Here we investigated the impact of these two subpopulations on the activity of the local CFA circuit as well as on the behavior in different contexts. When rats were not involved in a task, the effect of RFA was mainly excitatory in the deep CFA layers, while the superficial layers remained unaffected. This can be interpreted as a non-selective activation of the deep CFA neurons enabling a variety of spontaneous movements. During the preparation phase before a movement, the RFA had an opposite impact on the superficial and deep layers: while the superficial CFA layers were excited by RFA input, the deeper layers were mostly inhibited, minimizing movements and enabling continued holding of a lever. During the movement phase, the inhibitory effect on neurons in the deep CFA layers was counterbalanced by excitation, thus enabling a selected conduction of movements. The opposing effects during preparation and movement phase on CFA deep layers were correlated with increased firing rates of the RFA preparation and movement subpopulations, respectively, making it likely that the inhibition resulted from increased activities of these subpopulation specifically. With an electron microcopy approach we demonstrate that inhibitory and excitatory CFA neurons are directly targeted by RFA, thus providing a mechanism for the bidirectional control of CFA activity. Please note that the depicted impact of RFA on excitatory or inhibitory CFA neurons refers to net effects in this figure, not to the targeting of individual neurons.

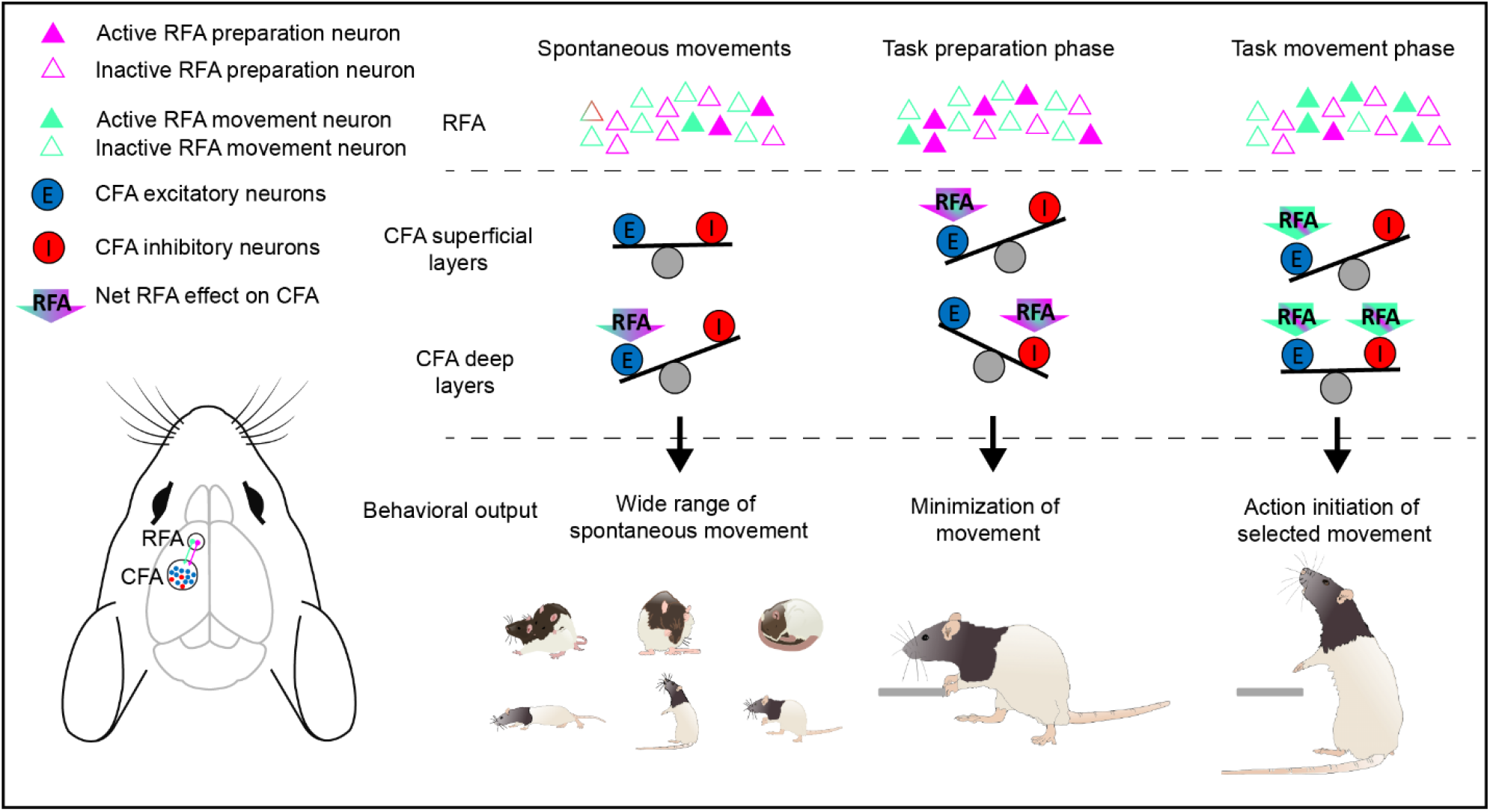

## Main

The control of movement is a central aspect of life, and understanding the neural basis of this process can provide insights into a wide range of phenomena related to movement planning and execution, including motor learning and skill acquisition, voluntary and involuntary movements, decision-making and goal-directed behavior. In rodents, the rostral and caudal forelimb areas (RFA and CFA) are thought to play important roles in the control of forelimb movements^5,6^, but the interaction between these areas during movement planning and execution are far away from being well understood. While the activity of RFA and CFA neurons is similar across a range of tasks^7–9^, it remains unclear which information is conveyed by neurons projecting from RFA to CFA during different phases of motor planning and execution, and how these projecting neurons shape CFA activity during these different epochs.

## Results

### Freely moving rats responded to a vibro-tactile delayed go-cue

To investigate the role of RFA and CFA in movement preparation and execution, we developed a preparation-movement task for freely moving rats^10,11^. Rats initiated trials by pulling a lever with their forepaw. They then had to keep holding the lever until a vibrotactile stimulus (0.3 s), which served as the go cue, was delivered via the lever (Fig. 1a). To discourage timing, rats were pseudo-randomly required to hold the lever for 0.6 s or 1.6 s (short or long delay; Fig. 1b).

**Fig. 1:**
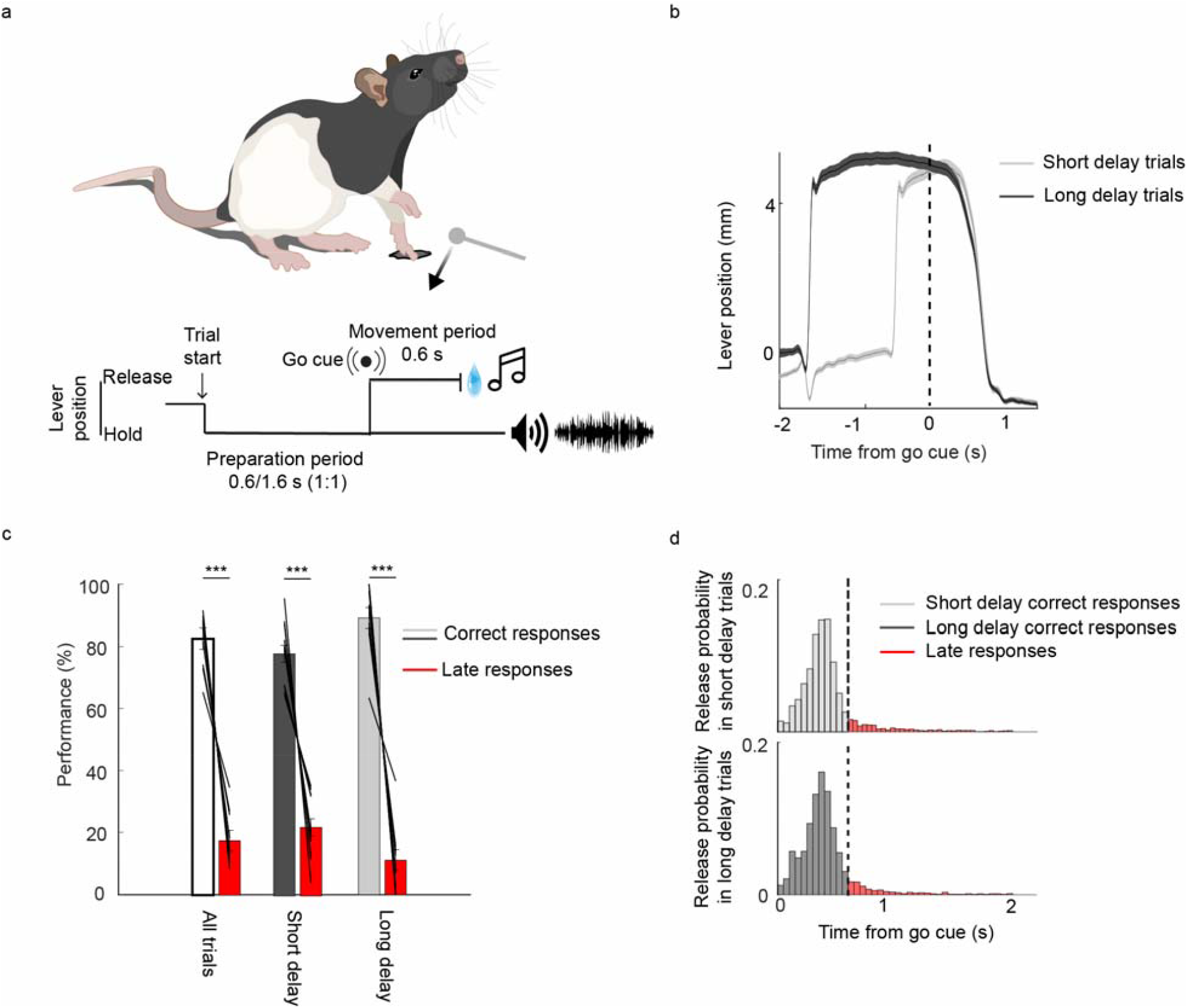
Preparation-movement task. **a**. Simplified illustration of the task. To initiate a trial, the freely moving rat had to move a lever horizontally and hold it. After a preparation period of 0.6 or 1.6 s, the lever vibrated, which was the go cue. Following the go cue, the rat had to release the lever within a response window of 0.6 s to receive a reward; otherwise, the trial was considered a late trial. Releases during the preparation period were considered early trials. Correct trials were signaled with a click tone and the late trials were followed by a 1 s timeout and white noise. **b**. Rats learned to steadily hold the lever during the preparation period and to release the lever after the go cue. **c**. Behavioral performance in the last training session of each animal (n = 11 rats, n = 114 sessions; correct responses in all trials: 82.6%, correct responses in trials with short delay: 78.0%, correct responses in trials with long delay: 89%). **d**. Reaction time distributions display a prominent peak after the go cue in both short (top) (N = 11 rats, n = 114 sessions, n = 4,159 trials) and long (bottom) delay trials (N = 11 rats, n_short_ = 4,159 trials, n_long_ = 3,825 trials), indicating that the rats attended to the cue. ***p<0.001, paired t-test; error bars: SEM. See also Extended Data Table 1.

After the go cue, the rats had to release the lever within a response window of 0.6 s to receive a water reward (Fig. 1a). If the rats released the lever before the cue or after the response window (0.6 s), the trial was considered an error trial and classified as early or late, respectively. After training, rats responded correctly to the go cue in more than 70% of the trials with both short and long delays (short delay trials: 78.2 +/-3.1 %, long delay trials: 88.1 +/-3.2%, all data are presented as the mean □ ± □ standard error of the mean (SEM) in the rest of the manuscript unless otherwise noted, p=0.029, two sided paired t-test, n = 11 rats; Fig. 1c). The higher performance in long trials emphasized the importance of the hold period for the preparation of the movement. The reaction times (RT) distributions had a sharp peak after the go cue onset in both the short and long delay trials, confirming that the rats responded to the cue (Fig. 1d, Extended Data table 1). Taken together, the rats learned our preparation-movement task well. Importantly, this task entails distinct epochs for movement preparation (hold) and movement execution (release) and is therefore ideally suited to disentangle the contributions of RFA and CFA to movement preparation and execution.

### Preparation neurons are more common in RFA than CFA

To characterize neuronal responses relating to movement preparation and execution across motor cortical areas during the task, we implanted 32-electrode laminar silicon probes in the RFA or CFA of ten rats split into two groups of five per forelimb area (Fig. 2a, Extended Data table 2). We recorded 240 neurons in the RFA and 355 neurons in the CFA (3–4 sessions per rat). Depending on the electrode depth^12^, we classified neurons above and below 0.75 mm as superficial or deep, respectively. In rats, the superficial layers have the same thickness in premotor and primary motor areas, with the premotor areas characterized by a thicker layer 6^13^. The vast majority of the neurons were modulated by the task (228/240 (95%) in the RFA, and 336/355 (95%) in the CFA); minimum of three bins with an absolute z score higher than 1.96, corresponding to 95% confidence intervals).

**Fig. 2:**
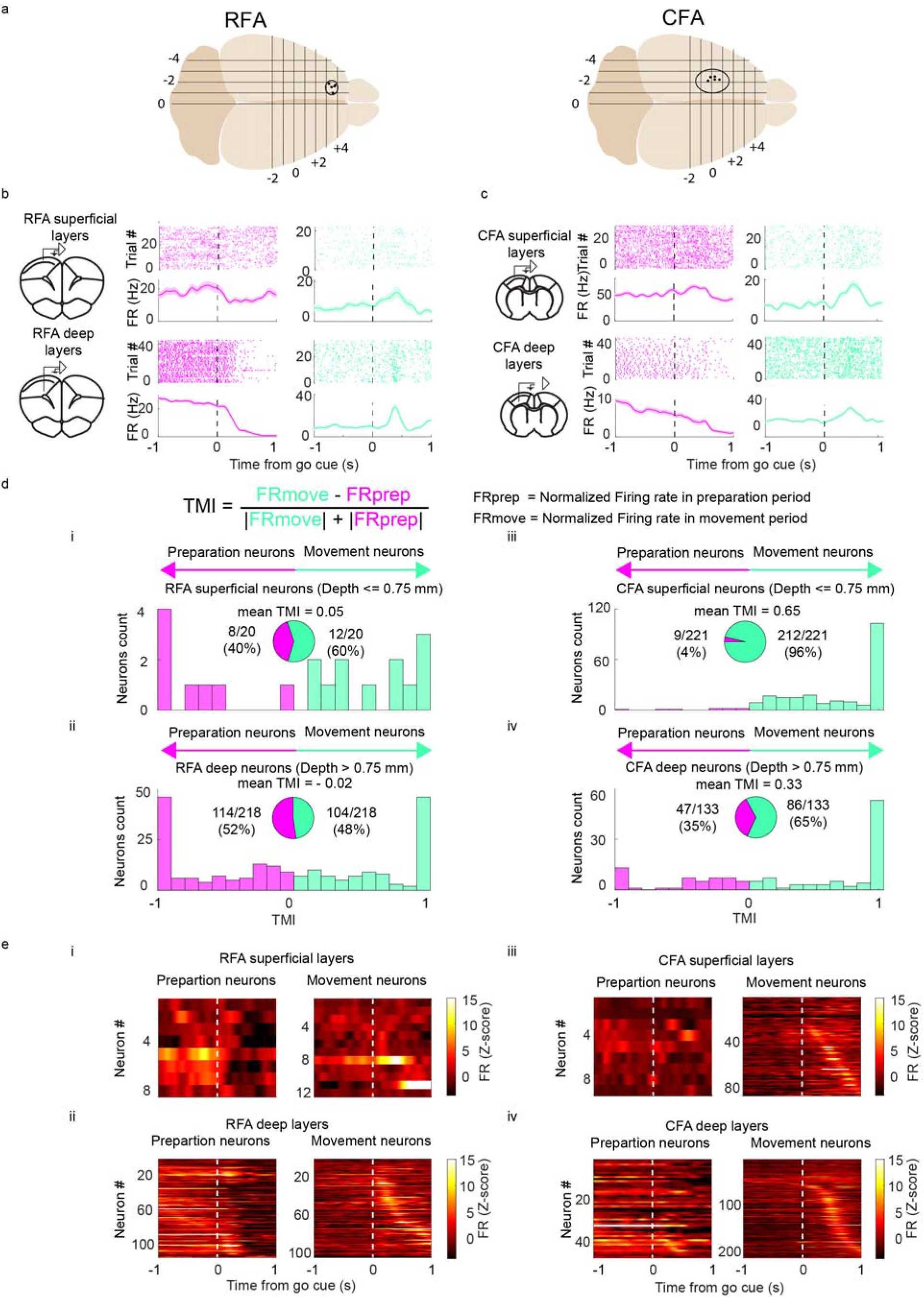
Activities of neuronal subpopulations in RFA and CFA. **a**. Reconstructed laminar silicon probes implantation sites in the RFA and CFA (N = 5 rats for each area). **b**. Raster and peristimulus time histogram (PSTH) of example neurons in RFA superficial (top) and deep layers (bottom). Neurons modulated during preparation (magenta) and movement (cyan) periods were observed in both superficial and deep layers. **c**. Same as b but in the CFA. **d**. Neurons were classified based on their mean firing rates during the preparation and movement periods. If a neuron had a positive task modulation index (TMI), it was classified as a movement neuron, otherwise, it was classified as a preparation neuron. The RFA contained similar proportions of preparation and movement neurons (i and ii). In contrast, the CFA had a strong bias towards movement neurons (iii and iv). **e**. Normalized firing rates (Z-score; methods) of all modulated neurons in RFA superficial layers (i), RFA deep layers (ii), CFA superficial layers (iii), and CFA deep layers (iv). dashed lines: go cue; magenta: preparation neurons, cyan: movement neurons; The shaded background shows SEM. Coronal sections in b and c adapted from^77^. See also Extended Data Fig. 1 and table 2.

Neurons were predominantly active during either the preparation or movement period (Figs. 2b,c). To classify neurons as either preparation or movement neurons, we used a task modulation index (TMI) (TMI = (FRmove – FRprep)/(|FRmove| + |FRprep|), where FR is the normalized z-score of the firing rate in the respective period. A positive TMI identified a movement neuron, while a negative TMI identified a preparation neuron (Fig. 2d). In both superficial and deep layers, we found a higher proportion of preparation neurons in the RFA than in the CFA (Fig. 2d). The RFA contained similar proportions of preparation and movement neurons, in line with a role of RFA in motor preparation^14,15^, and similar to other premotor areas in rodents^3,16–20^, consolidating the homology of the RFA and primate premotor areas^6,21^. The TMI distribution for the RFA had no particular bias for preparation or movement (mean RFA superficial neurons TMI = 0.06 +/-0.17, mean RFA deep neurons TMI = -0.02 +/-0.05; Fig. 2d, Extended Data Fig. 1a). In contrast, most CFA neurons had higher firing rates during the movement period, meaning that the TMI distribution had a clear bias towards activity during the movement period (mean CFA superficial neurons TMI = 0.65 +/-0.03, mean CFA deep neurons TMI = 0.33 +/-0.06; Fig. 2d, Extended Data Fig. 1b) and suggesting that CFA is primarily involved in movement execution. Overall, both RFA (Fig. 2e i, ii) and CFA (Fig. 2e iii, iv) were modulated by the task with similar activity patterns albeit with different proportions.

### The temporal hierarchy runs from RFA to CFA

The differential distribution of preparation and movement neurons across RFA and CFA suggests a functional hierarchy between the two regions. We therefore asked whether this hierarchy is reflected in the timing of neuronal activity, with the hypothesis that RFA precedes CFA, similar to the neuronal activity in a reaching task^22^ and reflecting top-down RFA to CFA connectivity^6,13,23^. Because of the small number of superficial neurons in the RFA and superficial preparation neurons in the CFA (Fig. 2d i, ii), we pooled superficial and deep datasets (note that all results described below also remain valid when only analyzing the deep layers; Fig. 3, Extended Data Fig. 2, Table 3).

**Fig. 3:**
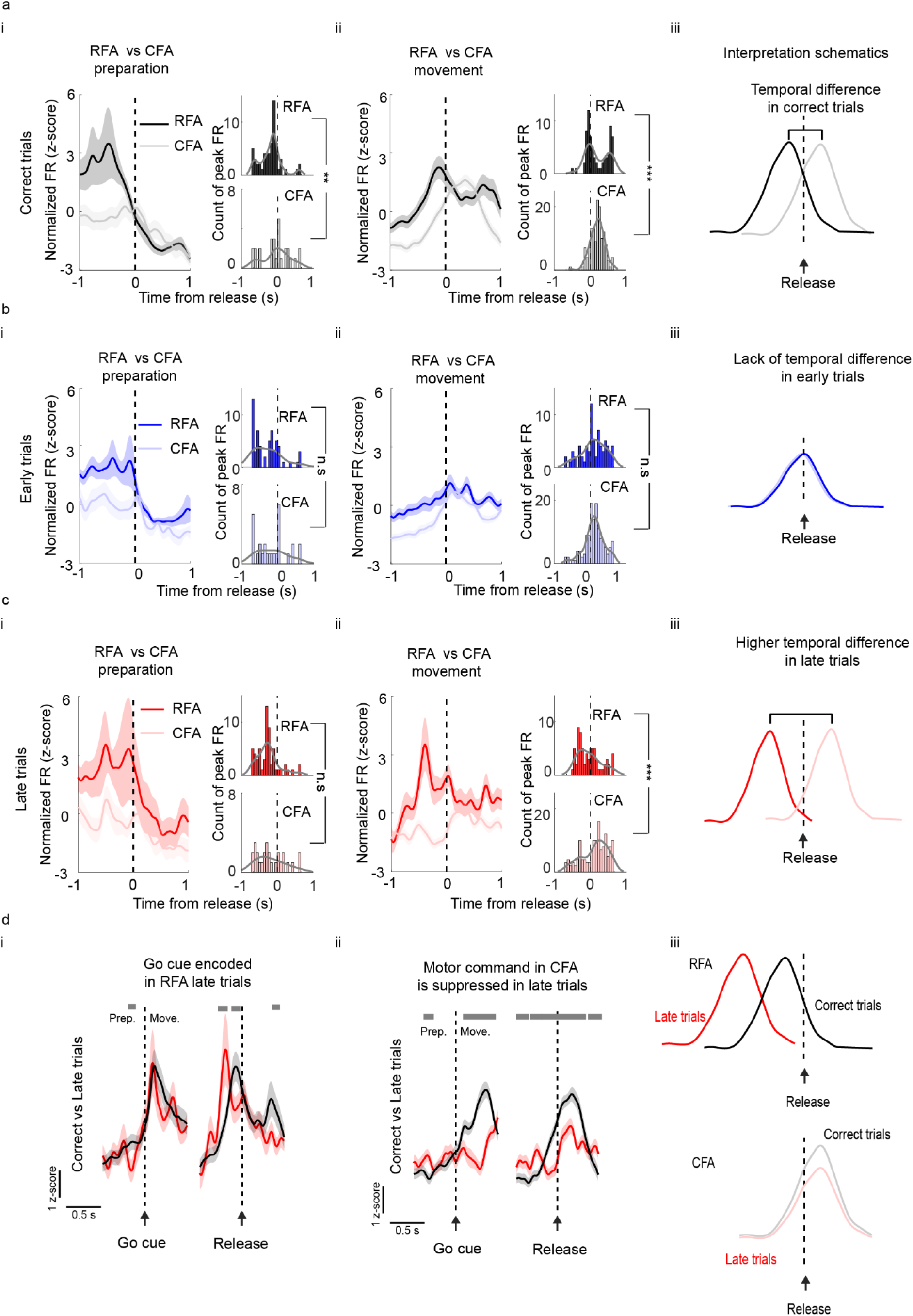
Firing rate peaks indicate a temporal hierarchy from RFA to CFA subpopulations. **a**. Normalized firing rates (z-score) and population distribution of peak activity during RFA and CFA preparation (i) and movement (ii) subpopulations in correct trials aligned to lever release. The activity of RFA subpopulations precedes those in the CFA in correct trials (iii). **b** and **c**. Same as **a** but for early and late trials, respectively. In contrast to correct trials, the temporal hierarchy is altered in error trials with no temporal difference in early trials and a more pronounced temporal separation in late trials. **d**. Average population activity of the movement subpopulations in the RFA and CFA in correct and late trials. Mean normalized firing rate (z-score) of RFA movement subpopulation aligned to go cue (i, left) and release (i, right). Mean normalized firing rate (z-score) of CFA movement subpopulation (ii). The go cue is encoded in the RFA movement subpopulation, and the motor command is encoded in the CFA movement subpopulation (iii). Blue: early trials, black: correct trials, red: late trials; the dashed black line refers to the go cue or release onset. Two-sample Kolmogorov-Smirnov tests were used for peak distributions in **a–c**. Two-way ANOVAs were used in **d** for repeated measures; the top gray bar indicates significant differences (p<0.05). The shaded background shows SEM. See also Extended Data Fig. 2,3 and table 3.

In line with our hypothesis, the activity of the preparation subpopulation peaked significantly earlier in the RFA than in the CFA relative to movement execution in correct trials (RFA_prep_ correct trials: - 0.30 +/-0.06 s, CFA_prep_ correct trials: - 0.05 +/-0.10 s; N_RFA_ = 5 rats, n_RFA-prep_ = 58 neurons, N_CFA_ = 5 rats, n_CFA-prep_ = 27 neurons; p = 0.005, two-sample Kolmogorov-Smirnov test; Fig. 3a i). Activity in the movement subpopulations also peaked earlier in the RFA than in the CFA, although the distribution of peak firing rates in the RFA was bimodal, with some neurons preceding and some lagging CFA activity (RFA_move1_: - 0.06 +/-0.03 s, CFA_move_: 0.29 +/-0.02 s, RFA_move2_: 0.77 +/-0.03 s; N_RFA_ = 5 rats, n_RFA-move_ = 71 neurons, N_CFA_ = 5 rats, n_CFA-move_ = 135 neurons; p = 4.9*10^−4^, two-sample Kolmogorov-Smirnov test; Fig. 3a ii). Taken together, the interareal timing differences of both preparation and movement subpopulations support the view of a hierarchical information flow from RFA to CFA (Fig. 3a iii). This timing was not conserved in error trials, with the temporal delay between RFA and CFA depending on the type of error (Figs. 3b, c). In early trials, preparation neurons in RFA and CFA were recruited almost simultaneously (Fig. 3b i). In addition, the movement subpopulations in RFA and CFA both peaked after lever release (Fig. 3b ii). In contrast, we found the temporal delays between RFA and CFA subpopulations increased in late trials (Fig. 3c i-iii). These shifts in peak neuronal activity in the RFA and CFA offer a glimpse into the RFA to CFA communication which seemed to break down in error trials.

### Temporal delays in late trials originate in the movement subpopulation of the CFA

To investigate the transition between preparation and movement in both forelimb areas in more detail, we aligned the firing rates of correct and late trials to the go cue and release time. This allows testing if neuronal responses were time-locked to the cue or the movement. Here, we focused on the movement subpopulations because they are the ones with a direct correlation with the go cue as well as movement. When aligning the RFA movement subpopulation to the go cue, correct and late trials followed similar trajectories (Fig. 3d i); both trial types were associated with a sharp increase in neuronal activity after the go cue (Extended Data Fig. 3). The similar neuronal responses in both trial types indicated a correct detection of the go cue in the RFA even in late trials; therefore, the transfer into a motor command must have been delayed elsewhere. In contrast, the CFA movement subpopulation did not respond with the same latency relative to the cue in correct and late trials (Fig. 3d ii). Instead, late trials were characterized by a drop in neuronal activity after the go cue when aligned to the release time; the neuronal responses in CFA became time locked for both correct and late trials. However, late trials were characterized by an overall lower neuronal activity. The decreased neuronal activity was already apparent before the release, indicating that the CFA’s neuronal state in late trials might have been suboptimal. Taken together, the go cue was correctly encoded in the RFA in both correct and late trials, but the putatively suboptimal neuronal state in the CFA in late trials might have delayed the motor command (Fig. 3d iii).

### Neurons projecting from RFA to CFA are involved in movement preparation and execution

Given that RFA activity precedes CFA activity in correct trials and that the course of delayed movements might have their origin in the CFA but not in the RFA, we hypothesized that the RFA predominantly conveys information about the planning of the movement but does not provide the actual execution signal to the CFA. To test this hypothesis, we asked how identified subpopulations in the RFA communicate with the CFA, and how they affect behavioral performance. To identify RFA neurons projecting to the CFA, we employed optogenetics to induce antidromic spikes in the RFA by stimulating RFA axons in the CFA (Fig. 4a). We injected AAV5-hSyn-ChR2-eYFP into the RFA of three rats, implanted a laminar silicon probe in RFA and inserted an optical fiber into CFA (Figs. 4a–c, Extended Data Table 4; Methods). We found that both preparation and movement neurons project from RFA to CFA (Figs. 4 d, e; Extended Data Fig. 4). In total, we identified 21 out of 46 neurons (46%) projecting from RFA to CFA. Out of the 21 identified projecting neurons, 11 (52%) were primarily active during the preparation period (negative TMI) and the rest of the neurons were mainly active during the movement period (positive TMI, 10/21, 48%; Fig. 4f). Thus, neurons projecting from RFA to CFA convey both planning- and movement-related information to the CFA.

**Fig. 4:**
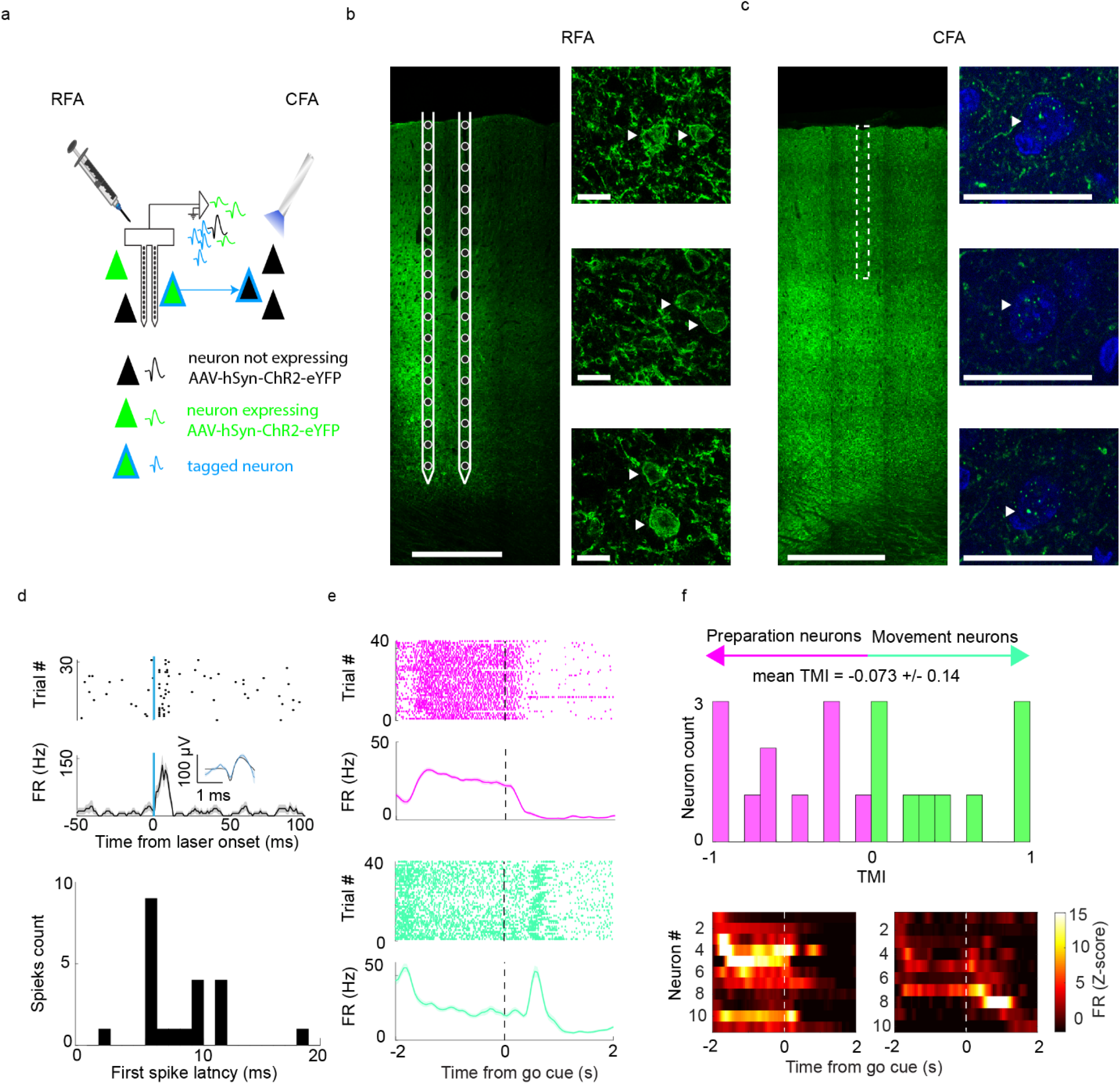
RFA to CFA projection conveys both preparation and movement information. **A**. Schematic of the phototagging experiment. The RFAs of rats (N = 3) were injected with AAV5-hSyn-ChR2-eYFP. A laminar silicon probe was also implanted in RFA and an optical fiber was emplaced in their CFAs. At the end of the session, each rat was stimulated with 1 ms light pulses delivered to their CFA via the optical fiber. The light pulses induced antidromic spikes by activating ChR2 in RFA axonal terminals in CFA (light blue waveforms). **B**. Coronal section through the RFA with cells expressing AAV5-hSyn-ChR2-eYFP; white circles: estimated electrode positions in RFA; scale bar = 500 µm (left). Examples of RFA neurons expressing AAV5-hSyn-ChR2-eYFP; scale bar 20 = µm (right). **c**. Coronal section of the CFA with RFA axons expressing eYFP in CFA; dashed white rectangle, optical fiber position in CFA; scale bar = 500 µm (left). Examples of CFA neurons labelled with DAPI (blue) close to RFA axons; scale bar = 5 µm (right). **d**. Raster and PSTH of a tagged neuron projecting from RFA to CFA (top). Inset: mean waveform of simultaneous spikes (black) and antidromically-induced spikes (light blue). Latency of the first spike after the light pulse (Bottom). **e**. Raster and PSTH of an example preparation (top) and movement neuron (bottom) projecting from RFA to CFA. **f**. Task modulation index (TMI) for RFA neurons projecting to the CFA (top). **F**. Normalized firing rates (z-score) of all preparation (bottom left, N = 3 rats, n = 11 neurons) and movement neurons (bottom right, N = 3 rats, n = 10 neurons) projecting from RFA to CFA. The shaded background shows SEM. See also Extended Data Fig. 4 and table 4.

### Inverse reinforcement learning predicts opposing roles of projecting preparation and movement subpopulations

We were puzzled by the similar percentages of preparation and movement neurons projecting from RFA to CFA, which did not match our initial hypothesis that the RFA mainly conveys a planning signal to CFA. Because we do not have the means to individually manipulate one of these functional subpopulations with optogenetics, we examined whether the projecting preparation and movement subpopulations might contribute differentially to the task through the lens of an inverse reinforcement learning algorithm termed NeuRL^24^. Reinforcement learning is a (mathematical) framework that formalizes the optimization of collected reward from an agent (in this case a rat) which applies actions (e.g. hold or release) in states (here defined by temporal bins) in an environment upon solving a task. Inverse reinforcement learning (as performed by NeuRL) then turns the optimization procedure upside down in that the intrinsic immediate reward function of an agent has to be inferred from recorded behavior. By mapping neuronal activity onto the generated reward, NeuRL can be leveraged to predict the behavioral effects of selective manipulation of functionally (or arbitrarily) defined subpopulations with simulated manipulations. Here we used NeuRL to simulate the inhibition of different subpopulations in RFA.

We used the lever release times (behavior) and the activity of RFA neurons as inputs to NeuRL (Fig. 5a, Extended Data Fig. 5a). We extracted a reward estimate via inverse reinforcement learning from the behavioral data and mapped features (i.e., mean FR in 0.2 s time bins) derived from the activity of recorded RFA neurons onto these rewards. Through this mapping, we obtained reconstructed reward values for each RFA neuron contributing to the reward with different weights (feature matrix; Fig. 5b, Extended Data Fig. 5b). Using this feature matrix, we simulated perturbations to different subpopulations in the RFA. With these simulated inhibitions, we computed perturbed reward values and subsequently used them to predict new actions via regular, forward reinforcement learning (Fig. 5c, Extended Data Fig. 5c; methods).

**Fig. 5:**
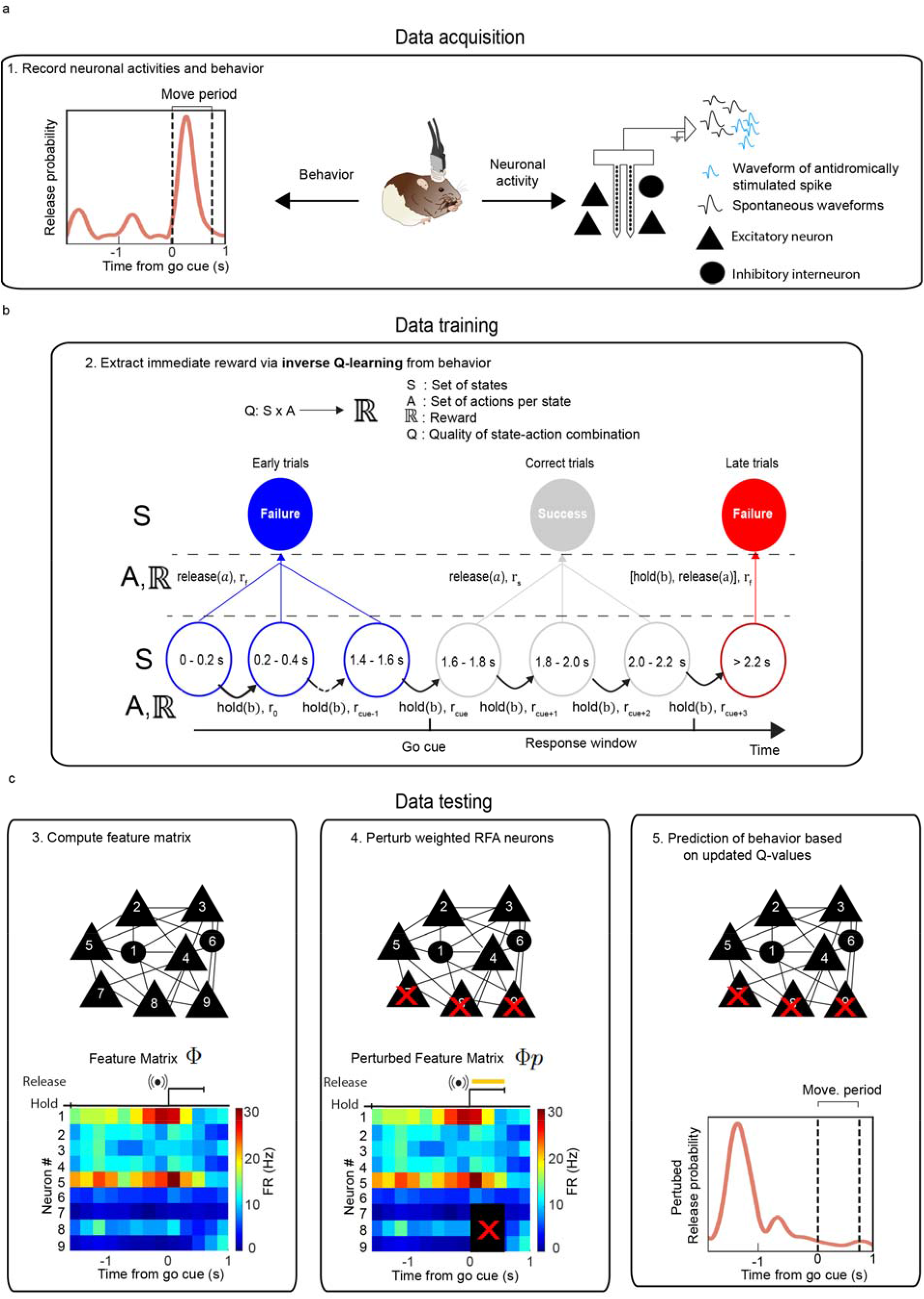
Q-learning workflow. **a**. The inverse Q-learning framework used the rats’ behavior (lever release times) and neuronal activity (1), consisting of the mean firing rates of RFA neurons in single trials, to assign functions to different neuronal subpopulations (N = 3 rats). Recordings from neurons identified as projecting from RFA to CFA were used to predict the trial outcomes and RTs. **b**. Training the network: a reward value was estimated from actions (hold referring to action b and release referring to action a) and states (success or failure) over 0.2 s bins for each trial. During the preparation period (0 to 1.6 s; blue circles), a state was termed ‘success’ if the action was hold and ‘failure’ when the action was release (2). In the movement period, a state was termed ‘success’ if the action was release, and after the movement period, a state was termed ‘failure’ regardless of the action. Based on the actions and states, a reward value was extracted via inverse Q-learning. This reward value was mapped onto a feature matrix reconstructed from the mean firing rate for each RFA neuron in 200 ms bins, giving each neuron a weight that contributed to the computed reward. **c**. Testing the network: based on the extracted rewards, different weights were assigned to the neurons in the behavioral task (3). Perturbation of different neurons in the feature matrix (4) produced a new perturbed reward value that was computed via forward Q-learning and then used to predict the perturbed behavior (5). For detailed mathematical equations see Methods and Extended Data Fig. 5.

To investigate the contributions of individual neurons to motor preparation and execution, we perturbed the neuronal activity at the end of the preparation period (0.5 s before the go cue) or the start of the movement period (starts with time of the go cue until 0.5 s after the go cue, covering most of the RTs distribution; Fig. 1d, Fig. 6a). First, we simulated the selective inhibition of all neurons projecting from RFA to CFA (Fig. 6b). NeuRL predicted that inhibiting all such neurons during the preparation period would result in significantly less early trials while an inhibition during the movement period would result in significantly more late trials (blue bar: 16.6% less early trials upon inhibition in the preparation period, unpaired t test, p = 1.3*10^−111^; red bar: 9.3 % more late trials upon inhibition in the movement period, unpaired t test, p = 1.2*10^−47^; Fig. 6b left). In accordance with these changes, inhibition during the preparation period predicted significantly shorter RTs and inhibition during the movement period resulted in significantly longer RTs (15 ms shorter after inhibition during the preparation period, unpaired t test, p = 0.003, 70 ms longer after inhibition during the movement period, unpaired t test, p = 4.6*10^−51^; Fig. 6b right).

**Fig. 6:**
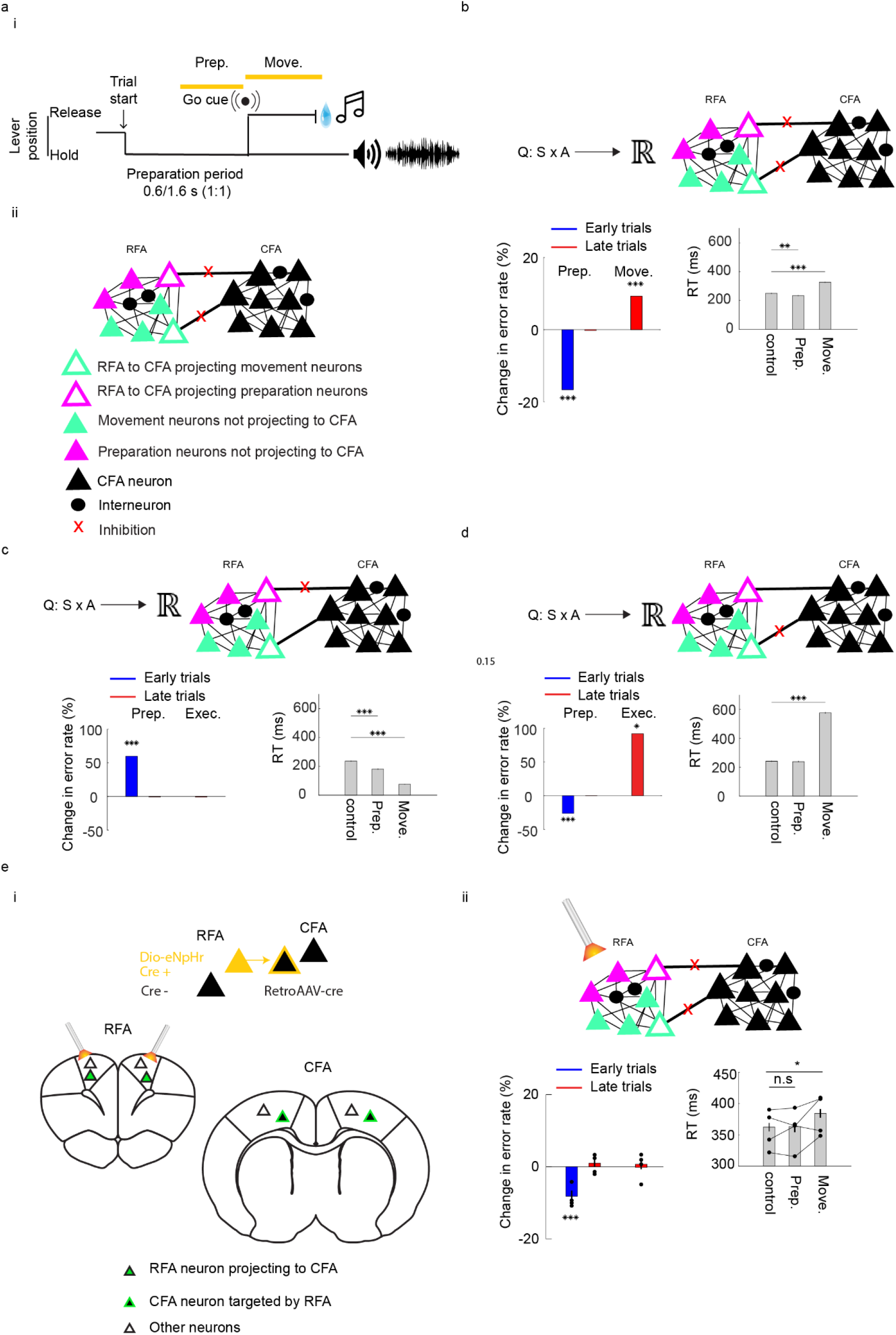
RFA to CFA-projecting neurons promote action but the two projecting subpopulations play opposing roles. **a**. Schematic for the inhibition experiments. We inhibited RFA to CFA-projecting neurons during either the preparation or movement periods *in silico* (N = 3 rats, 1,000 simulations for each rat), using NeuRL, or *in vivo* (N = 4 rats) (i). Schematic illustration of the different subpopulations (ii). **b**. Effect of inhibiting all RFA to CFA-projecting neurons *in silico* (n = 21 neurons) during the preparation period on error rate and RT. Neuronal inhibition significantly decreased early trials (left) and significantly shortened RT while inhibition during the movement period significantly increased late trials and increased RT (right). **c**. Same metrics as in b but relating to inhibition of the RFA to CFA preparation subpopulation (n = 10 neurons) *in silico*. We predicted significantly more early trials upon inhibition during the preparation periods (left). Further, we predicted shorter RT when inhibiting during the preparation or the movement period (right). **d**. We predicted that inhibiting the RFA to CFA-projecting movement subpopulation (n = 10 neurons) *in silico* during the preparation period would cause significantly fewer early trials and more late trials when inhibited during the movement period (left). Further, we predicted longer RTs when inhibiting during the movement period (right). **e**. Validating Q-learning with optogenetics. AAV-D-eNpHr-eYPF was injected bilaterally into the RFA and a retrograde viral vector carrying cre recombinase was injected into the CFA (i). Optical fibers were implanted bilaterally into the RFA. Inhibiting all RFA to CFA-projecting neurons significantly decreased early trials when inhibiting during the preparation period and significantly increased RT during the movement period (ii). Separated lines represent individual animals; * p<0.05, **p<0.01, *** p<0.001, unpaired t-test for NeuRL simulations (b–d), repeated measure two-way ANOVA for optogenetics (e). Coronal sections in e (i) adapted from^77^. See also Extended Data table 5.

The NeuRL approach allowed further discrimination between the projecting preparation and movement subpopulations. In line with previous findings in motor cortex^25^, we found two competing subpopulations with counteracting roles. Inhibiting the preparation subpopulation during the preparation period induced significantly more early trials and significantly shorter RTs (60% more early trials, unpaired t-test, p = 1.0*10^−85^ and 55 ms shorter RTs, p = 3.3*10^−6^; Fig. 6c). Inhibition of the same subpopulation during the movement period caused 0.7% fewer late trials (not significant) and 160 ms shorter RTs (p = 3.9*10^−270^). In contrast, inhibition of the movement subpopulation during the preparation period resulted in significantly fewer early trials but no significant effect on RTs (26% fewer early releases, p = 8.3*10^−264^; 2.5 ms shorter RTs; Fig. 6d), while the inhibition during the movement period resulted in significantly more late trials and significantly longer RTs (92% more late trials, p = 0.014; and 336 ms longer RTs p = 4.9*10^−186^; Fig. 6d). In other words, NeuRL assigned the preparation subpopulation a role in action suppression while the movement subpopulation was associated with action promotion.

### *In vivo* optogenetic inhibition confirms the role of neurons projecting from RFA to CFA predicted by NeuRL

We next aimed to confirm that NeuRL provided biologically-plausible hypotheses. Because of the technical limitations of currently available optogenetic tools, we focus here on the first hypothesis about the effect of inhibiting all neurons projecting from RFA to CFA. We injected the cre-dependent viral vector AAV5-hSyn-DIO-NpHR-EYFP and the retrogradely travelling cre-carrying vector retroAAV-cre into the RFA and CFA, respectively, of four rats to express inhibitory opsins in neurons projecting from RFA to CFA (Fig. 6e i, Extended Data Table 5; Methods). In line with the NeuRL predictions, inhibiting the neurons projecting from RFA to CFA during the preparation period significantly affected rat performance by increasing the number of early trials (early trials increased by 8.2 +/-0.15 %; N = 4 rats, n = 10 sessions per animal; p = 8.3*10^−5^, ANOVA for repeated measures; Fig. 6e ii left) but had no significant effect on the RTs of correct trials (Fig. 6e ii right). During the movement period, inhibiting neurons projecting from RFA to CFA had no effect on performance but significantly increased RTs in correct trials (RT in no laser trials: 0.36 +/-0.006 s, RT in trials with laser on during the preparation period: 0.36 +/-0.009 s, RT in trials with laser on during the movement period: 0.38 +/-0.007 s; N = 4 rats, n = 10 sessions per animal; no laser versus laser during movement period p = 0.044, ANOVA for repeated measures; Fig. 6e ii). While the optogenetic experiment showed a less drastic impact on RT and error rate than the theoretical predictions (most likely due to biological variability), the results of the optogenetic experiments were still in line with the NeuRL predictions, validating the tool for theoretical predictions about the behavioral role of specific neurons.

### Optogenetic inhibition of neurons projecting from RFA to CFA during rest reveals a predominantly excitatory connection

So far, we have characterized the fraction of RFA to CFA projecting neurons, their response properties, and their impact on behavior. The missing piece remains the impact of these projections on CFA activity. To fill this knowledge gap, we optogenetically inhibited the neurons projecting from RFA to CFA during rest (i.e., the rats were not involved in a task) and simultaneously recorded CFA neurons with laminar probes spanning the entire cortical depth (Fig. 7a). Neurons projecting from RFA to CFA specifically modulated neurons in the deep layers of the CFA (neurons modulated in superficial layers: 0/16 (0%), neurons modulated in deep layers: 10/48 (21 %), which is consistent with previous anatomical findings^26^. Most modulated neurons decreased their activity, likely due to lack of excitatory input, and only a small fraction increased their activity (8/10 decreased activity, 2/10 increased activity; analysis windows: 0.5 s baseline period before laser compared to 0.5 s laser duration, paired t-test, p < 0.05; Fig. 7b). Interestingly, the two neurons that increased their activity (or were disinhibited) had narrow waveforms (Fig. 7a, inset), suggesting a role for CFA fast-spiking (FS) interneurons in this pathway. Overall, these results suggest that the RFA modulation results in an increased predominantly excitatory activity in CFA.

**Fig. 7:**
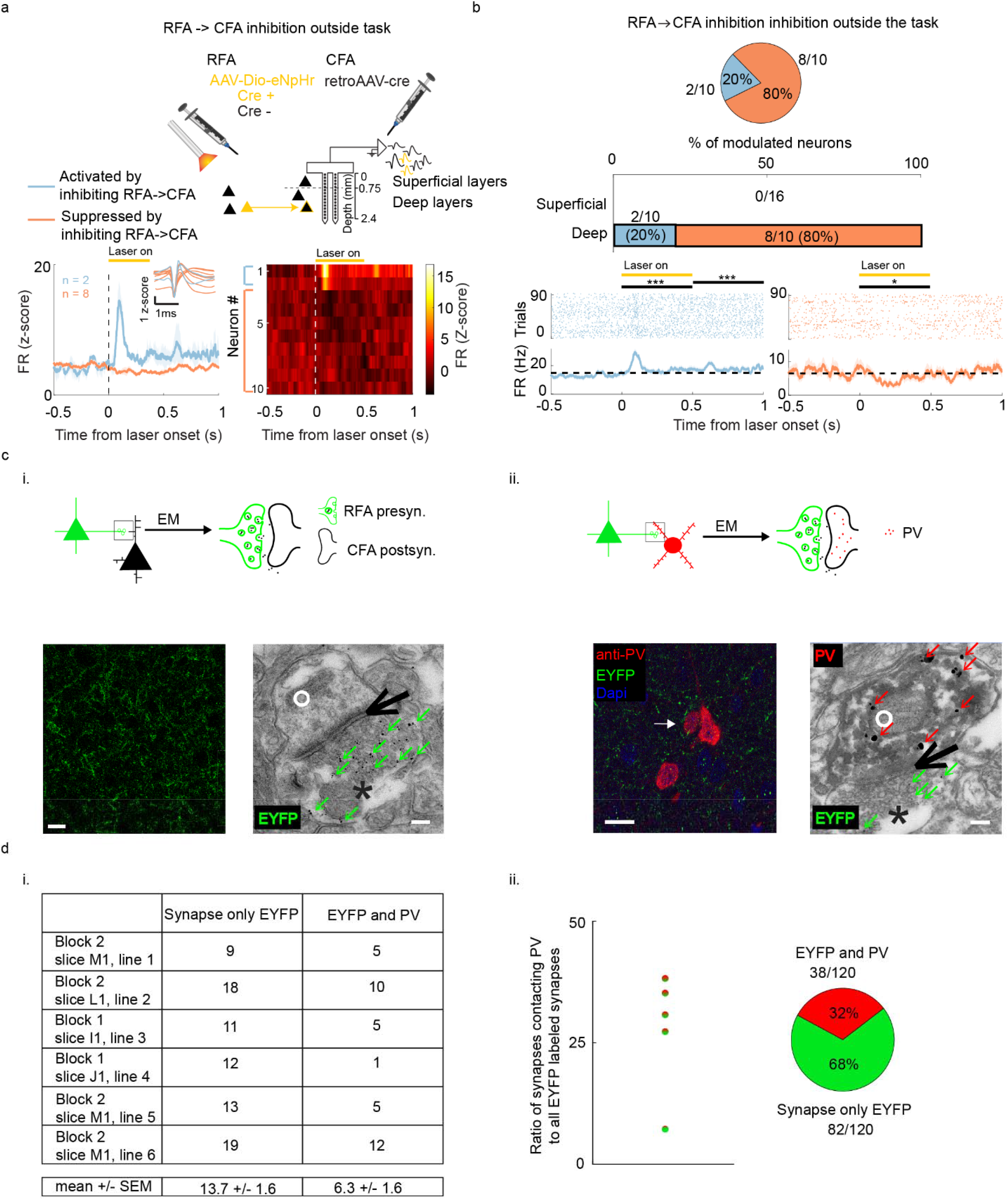
Neurons projecting from RFA to CFA modulate CFA neurons bidirectionally involving inhibitory interneurons. **a**. Schematic of injection and illumination scheme (upper panel). A subpopulation of neurons in the CFA (N = 3 rats, n = 10 neurons) was activated (light blue) while another was suppressed (light red) upon the inhibition of the RFA to CFA projection neurons outside the task (bottom left). Normalized firing rate (z-score) of all neurons modulated by the inhibition of neurons projecting from RFA to CFA (bottom right, one session each). **b**. The proportion of neurons in the superficial and deep layers activated and suppressed by inhibiting neurons projecting from RFA to CFA (top). Raster and PSTH example of a neuron activated and suppressed by inhibiting RFA to CFA projection neurons outside the task (bottom). **c**. RFA projects to excitatory as well as inhibitory neurons in CFA. Confocal image of coronal section through the CFA with AAV-DIO-eNpHr-eYFP expressing EYFP in RFA axons (i, left panel). Electron microscopy (EM) image in the CFA showing an RFA axonal terminal contacting a dendritic spine in CFA (i, right panel). Example of a PV interneuron in the CFA targeted by axons from RFA. Confocal image of a coronal section through the CFA expressing EYFP in RFA axons projecting to CFA. Neurons were labelled with an antibody against parvalbumin (PV). Confocal image example of coronal section (ii, left). EM image in the CFA showing an RFA axonal terminal contacting a spine of a PV+ interneuron in the CFA (ii, right). **d**. Counts of synapses with EYFP and synapses with EYFP and PV (i; N = 1 rat, n = 6 arbitrary lines). Ratio of RFA pre-synapses targeting PV+ interneuron post-synapses to total RFA EYFP+ pre-synapses in the CFA for different arbitrary lines (ii), note that line 1 and 2 have the same ratio. Scale bar in confocal images = 20 µm, scale bar in EM images = 100 nm; white arrow: PV+ interneurons; green arrows in EM: Immunogold staining against EYFP particles in RFA pre-synaptic terminal, red arrows: Immunogold staining against PV in CFA post-synapse, black asterisk: pre-synapse, black arrow: synaptic cleft, white circle: post-synapse; the shaded background shows SEM; * p < 0.05, *** p < 0.001, paired t test compared to 0.5 s baseline before laser onset. See also Extended Data Table. 5.

The disinhibitory effect of the projection specific optogenetic inhibition and the putative involvement of FS interneurons inspired us to test whether the anatomical connections can confirm the involvement of interneurons. In line with Dale’s principle^27^, the involvement of interneurons could be implemented in two ways: 1) Long-range GABAergic interneurons^28^ projecting from RFA to the CFA, or 2) RFA excitatory neurons projecting to CFA interneurons. To test these hypotheses, we used transmission electron microscopy to investigate the synapses onto neurons in the CFA (N = 3). Pre-embedding immunogold labeling of EYFP was used to identify EYFP labeled axonal terminals in contact with the dendritic spines of neurons in CFA (Fig. 7c). To test the first hypothesis, we classified synapses as symmetric (inhibitory) or asymmetric (excitatory). The vast majority of the synapses were asymmetric, ruling out the first hypothesis because asymmetric synapses originate from excitatory neurons (asymmetric synapses: 97/100, symmetric synapses: 3/100, Extended data Fig. 6a). Interestingly, RFA axons targeted dendrites across layers in the CFA (Extended data Fig. 6b), suggesting complex dendritic integration that is then transmitted to the soma. To test the second hypothesis, we stained coronal sections containing CFA with anti-PV antibodies and searched for EYFP-labelled axons across layers. Indeed, we found labelled axons that formed synapses onto PV+ dendrites (Fig. 7d i). In total, 32% (38/120) of the EYFP-labeled presynapses targeted PV+ interneurons (Fig. 7d ii). Considering that interneurons in the cortex only amount to ∼20% of the population and that ∼40% of these are PV+^29^, the ratio of RFA axons contacting PV+ synapses compared to other synapses implies that the RFA has a bias toward contacting PV+ interneurons in the CFA. In sum, RFA impacts the CFA local circuit by targeting both excitatory and inhibitory neurons with an unexpected bias towards PV+ interneurons.

### RFA to CFA projection modulates the CFA in a context- and layer-specific manner

Our electrophysiological findings so far point to a predominantly excitatory impact of RFA on CFA deep layers in rats not engaged in a task. However, it has been shown that the premotor cortex modulates motor cortex superficial layers in a context dependent manner^30,31^. Further, we found that the CFA receives information across layers (Extended data Fig. 6). Both movement and preparation neurons in RFA project to CFA and are preferentially active during different task epochs which might lead to spatiotemporally complex effects on CFA activity during the task. Hence, we asked whether the RFA would have an impact on the superficial layers of the CFA while the rats were engaged in the task and whether the RFA impact on the deep layers of the CFA would remain predominantly excitatory during the preparation-movement task. To address these questions, we employed the same optogenetic dual virus strategy with separate injections into the RFA and CFA (Fig. 8a). We found that 11/27 (41%) and 66/137(48%) of the recorded CFA preparation and movement neurons were modulated upon yellow laser light during the task, respectively (Fig. 8b). The inhibition periods were identical to the ones used before (i.e., 0.5 s before the go cue and 0.5 s after the go cue, see Fig. 6a). Due to the low number of modulated preparation neurons (Fig. 8b) and the fact that there were no systematic differences between the two populations (Extended Data Fig. 7), we pooled the two populations here after. Out of the modulated neurons, 30/77 (39%) were modulated by the laser exclusively in the preparation period, 34/77 (44%) by the laser during the movement period, and a minority of the neurons were modulated by the laser in both periods (13/77, 17%, Fig. 8c). In line with the observation that the projection from RFA to CFA targeted both excitatory and inhibitory neurons in the CFA^32,33^ (Fig. 7c,d), we found that a subpopulation of the modulated neurons decreased its activity while another subpopulation increased its activity upon optogenetic inhibition. Contrary to the effects of optogenetic inhibition during rest, both deep and superficial neurons were modulated when inhibiting the projection from the RFA during the task. The majority of the neurons which decreased their activity by the laser during the preparation period were located in the superficial layers, while the majority of the neurons that increased their activity were hosted in the deep layers (superficial modulated neurons: 14/30 (47%), deep modulated neurons 16/30 (53%), increased activity superficial: 3/14 (21%), decreased activity superficial: 11/14 (79%), increased activity deep: 10/16 (62%), decreased activity deep: 6/16 (38%); Fig. 8d). Upon optogenetic inhibition during the movement period, neurons that decreased their activity were predominantly present in the superficial layers, while neurons in deep layers showed a balanced proportion of increase and decreased activities (superficial modulated neurons: 13/34 (38%), deep modulated neurons 21/34 (62%); increased activity superficial: 1/13 (8%), decreased activity superficial: 12/13 (92%), increased activity deep: 9/21 (43%), decreased activity deep: 12/21 (57%); Fig. 8e).

**Fig. 8:**
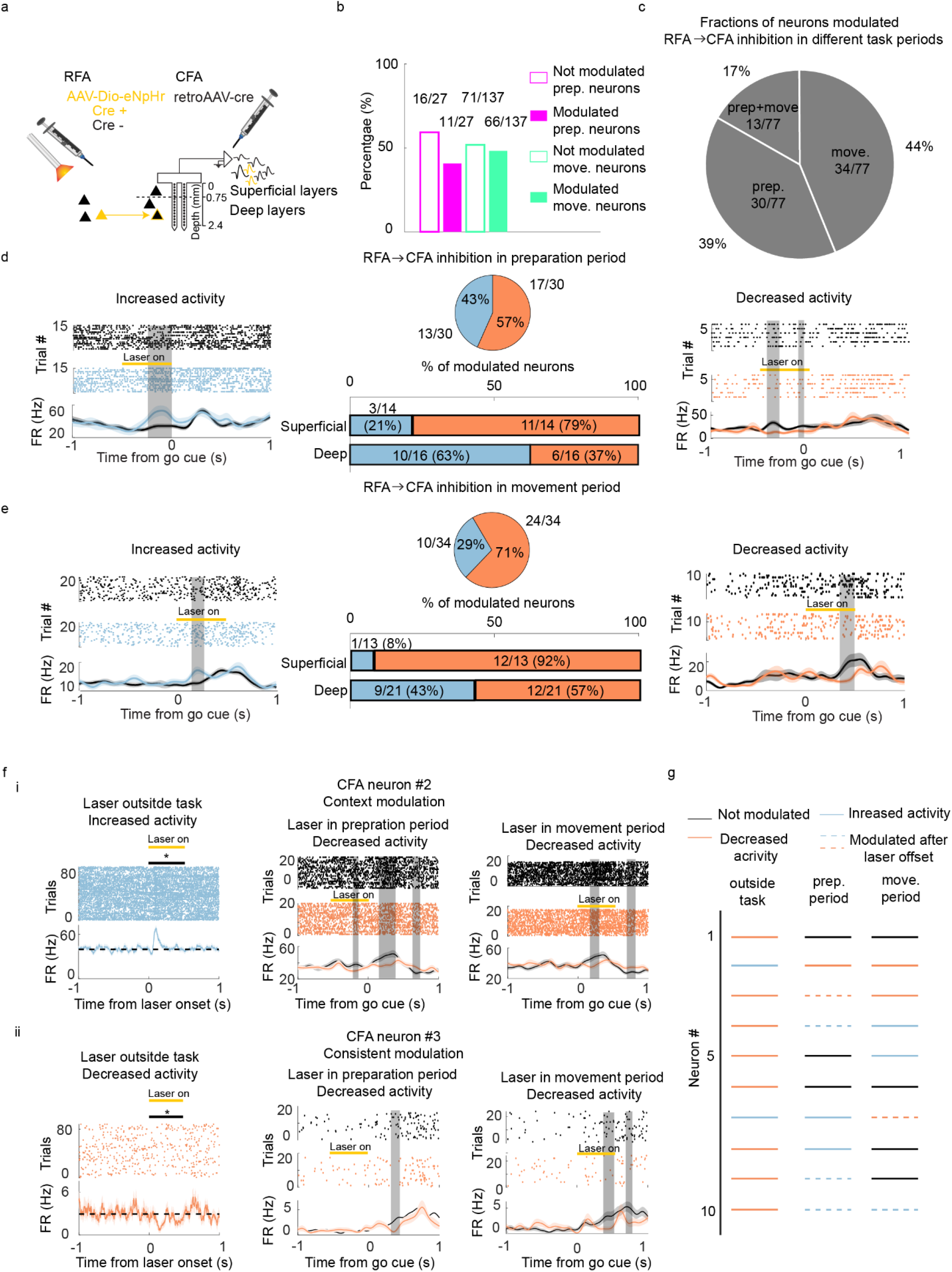
RFA to CFA projections affect CFA neuronal activity in a task phase and cortical layer dependent manner. **a**. Schematic of experimental strategy. AAV-Dio-eNpHr-eYFP was injected bilaterally into the RFA and a retrograde viral vector carrying cre recombinase (retroAAV-cre) was injected into the CFA. Optical fibers were implanted bilaterally into the RFA and a laminar silicon probe was implanted into the CFA (n= 3 rats). **b**. The proportion of CFA neurons significantly modulated by inhibiting RFA to CFA-projecting neurons. **c**. Fractions of CFA neurons modulated by optogenetic inhibition of RFA projection neurons during the different task periods. The task period refers to the laser on time. Neuronal modulation was tested during laser on. **d**. Effect of inhibiting RFA to CFA projection on CFA neurons. Example of modulated neurons that increased activity (left panel) and decreased activity (right panel) during the preparation period, respectively. The proportion of neurons that increased and decreased activity during the preparation period and their distribution in superficial and deep layers (middle panel). **e**. Same as **d** but when inhibiting the RFA to CFA projection during the movement period. Most modulated neurons decreased their activity during the movement period when inhibiting RFA to CFA projection. **f**. Neurons react differentially to the inhibition of RFA to CFA projection depending on whether the inhibition happened outside the task or during the preparation period or movement periods of the task. For instance, the activity of neuron #2 decreased by the manipulation outside the task (i, left panel), increased by the manipulation during the preparation period (I, middle panel), and remained unaffected by the manipulation during the movement period (I, right panel). In contrast, the activity of neuron #3 consistently decreased by the manipulation outside the preparation-movement task (i, left panel), as well as during the task (ii, middle and right panels). **g**. Summary of all neurons modulated outside the task and their response within the task (neurons # 1 to 10) (light blue bar: increased activity, light red bar: decreased activity, dashed bar: modulated after laser offset, black bar: not affected). Yellow bar: laser illumination. Gray shaded area: p<0.05, two-way ANOVA for repeated measure. Black: Trials without laser illumination, light red: neurons significantly decreased their activity by inhibiting RFA to CFA-projecting neurons, light blue, neurons significantly increased their activity by inhibiting RFA to CFA-projecting neurons, orange bar: time of laser illumination, grey bar: time bins significantly different than control; statistical tests were ANOVA for repeated measures for two curves comparison, gray shaded areas: p<0.05, paired t test for comparison to baseline outside the task, * p < 0.05; the shaded backgrounds shows SEM. See also Extended Data Fig. 7 and Table. 5.

In contrast to the dominant excitatory effect on the CFA outside the preparation-movement task, we found diverse effects during the task (Fig. 8f i, ii). For instance, a CFA neuron that increased its activity by optogenetic inhibition of the RFA projection outside the task decreased its activity by optogenetic inhibition of the same RFA neurons during both the preparation and movement period (neuron #2; Fig. 8f i), while another neuron responded similarly outside and within the task (neuron #3; Fig. 8f ii). The manipulation during the preparation period also led to a significant decrease in firing rate during the movement period (neurons #2 and #3; Fig. 8f i and ii, middle panels), indicating complex neuronal interactions during the task. Investigating this delayed impact of the optogenetic manipulation on a population level revealed that this was a common effect happening also in 76 neurons with no significantly modulated responses during laser light (Extended Data Fig.7). Outside the task, we observed this delayed effect in only one neuron which was also significantly modulated during the laser (see example neuron in Fig. 7b). These diverse effects suggest a more complex role for neurons projecting from RFA to CFA during the preparation-movement task than during rest (Fig. 8g).

To summarize, RFA projections to CFA reorganized subpopulations in the CFA in a dynamic and context-dependent manner, likely utilizing local inhibitory interneurons in CFA deep layers (Fig. 9). Please note that optogenetic inhibition removed the excitatory RFA input into CFA. Thus, the effects of the optogenetic inhibition have to be reversed to obtain the impact of the RFA input under physiological conditions. By doing so, we see that RFA inputs to the CFA had no impact on neurons in its superficial layers and were predominantly excitatory in the deep layers when the animal was not involved in a specific movement task. Conversely, when the animal engaged in a goal-directed task, with rich temporal patterns emerging during different task phases in the different subpopulations in the RFA, the impact of the RFA on neurons in CFA superficial layers was predominantly excitatory, and the impact on deep layers depended on the task phase. During movement preparation, RFA input to CFA neurons in deep layers was predominantly inhibitory, suppressing unwanted motor output. When the animal executed the movement, the RFA impact on CFA deep layers resulted in balanced excitation and inhibition, allowing the selection of the proper movement program.

**Fig. 9:**
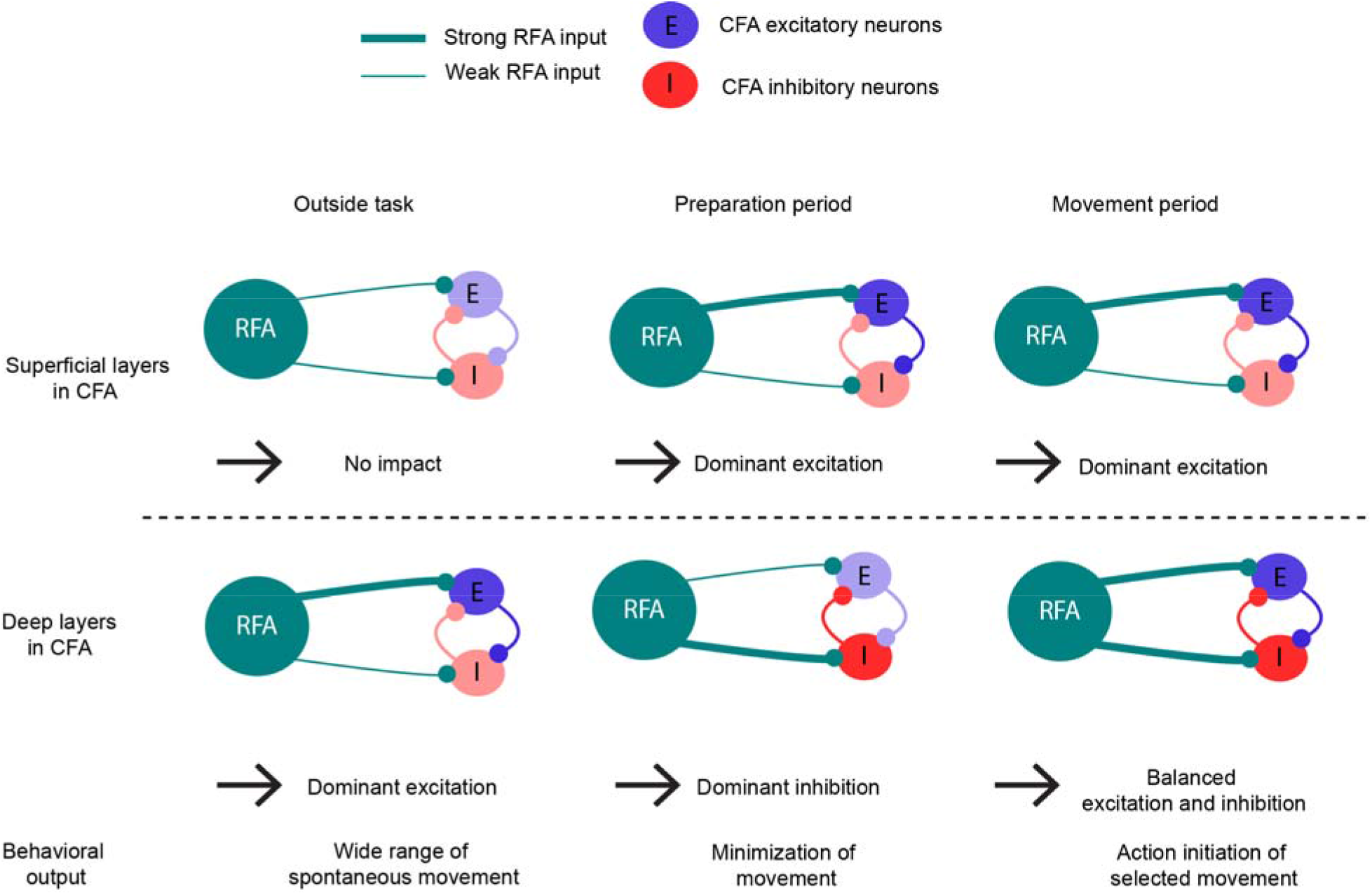
Schematic interpretation of the context- and layer-dependent impact of RFA on CFA. RFA (green circles) projection to CFA (green lines) impacts at least two different subpopulations in the CFA. The RFA targets both excitatory (blue circles) and inhibitory neurons (red circles) in the CFA. Depending on the layer and context, RFA input had different impacts on the CFA. The RFA did not impact CFA superficial layer activity during rest. During the preparation and movement periods, RFA input led to a net excitation in the CFA, amplifying CFA local circuits in the superficial layers. The RFA input to the CFA was mainly excitatory in its deep layers during rest, allowing a wide range of spontaneous movements. During the preparation period, the RFA input was predominantly inhibitory in the deep layers of CFA via the preferential activation of inhibitory neurons, and hence suppressed movements. During the movement period, the RFA bidirectionally modulated the CFA in the deep layers, leading to activation of one neuronal subpopulation and the suppression of another. Please note that the impact of RFA on excitatory or inhibitory CFA neurons refer to net effects, not to the targeting of individual neurons.

## Discussion

In this study, we characterized the neuronal activity in the RFA and CFA and the impact of RFA input on the CFA during different behavioral phases of a task to dissect the roles and interactions between these areas during preparation and movement execution. The work described here makes four distinct contributions to the field of motor neuroscience.

First, we confirmed that both RFA and CFA are involved in movement planning and execution and that the hierarchy flows from RFA to CFA as previously suggested^5–9,13–15,21,22,26,30,34–37^. In detail, we demonstrate here that the RFA contained a much higher number of preparation neurons than the CFA and that the activity of movement neurons in the RFA preceded their counterparts in the CFA. Importantly, we showed that this hierarchy was only present in correct trials, suggesting that the hierarchy is essential for performing a planned movement. Further, RFA neurons encoded the go cue in late trials, indicating that the signal to execute a movement was issued, but the motor command was delayed in the local CFA circuit. The movement subpopulation in the CFA was characterized by a lower firing rate in late trials, similar to its activity during the preparation period. The lower activity in CFA prior to movement in late trials could be a sign that the movement subpopulation was in a subspace that was suboptimal for generating movements^38,39^.

Second, with a phototagging approach, we revealed that the neuronal subpopulation from RFA to CFA contributes equally to planning and execution of movements. To go beyond this experimentally possible dissection, we developed an innovative machine learning-based tool (NeuRL) which allows the generation of predictions about neuronal subpopulations that are otherwise hard to assess due to their invariant responses to movement^40^. This novel machine-learning algorithm enables the functional dissociation of spatiotemporally-concurrent neuronal ensembles. To disentangle behavioral functions of the preparation and movement subpopulations, we combined NeuRL with electrophysiology and optogenetics. We identified neurons projecting from RFA to CFA with a phototagging technique and used reinforcement learning^41^ to assign functional roles to the preparation and movement subpopulations^24^. Analogous to the reward system in animals^42^, NeuRL finds an optimal strategy based on a feedback signal. While common decoding methods rely on supervised learning without accounting for the long-term consequences of the actions, NeuRL can explicitly assign a policy which maximizes reward over time. Reinforcement learning has recently been used to better understand neuronal activity and how it influences behavior^41,43^. However, the feedback signal is generally unknown. In contrast to previous approaches, we thus employed inverse reinforcement learning, which extracted reward values from the rat behavior instead of regular forward reinforcement learning, which assumes the feedback signal to be known a priori. This approach allowed us to assign functional meaning to neuronal subpopulations by conducting simulated inhibition studies based on a modified reward function inferred from the perturbed features. Thereby, we identified opposing roles of the preparation and movement subpopulations projecting from RFA to CFA with the first holding back movements and the second promoting actions.

Third, using immunohistochemistry and electron microscopy we identified a surprisingly strong role of inhibitory interneurons in the connection between premotor and motor cortex. Specifically, we found that RFA neurons project to both excitatory neurons as well as PV+ interneurons. Previously, it has been shown that RFA neurons project to fast-spiking PV+ interneurons in CFA^32^, and that these interneurons are activated prior to the forelimb reaching a target^44^, playing a role in shaping motor command^45^. Inputs to PV+ interneurons can be quite powerful as a single unitary excitatory postsynaptic potentials can evoke precisely timed action potentials in this cell type^46^. Here we added to this knowledge that RFA neurons are one of the sources that increase the activity of PV+ interneurons in CFA. In combination with the high activity of the preparation subpopulation projecting from RFA to CFA (Fig. 4e), the relatively low firing rate in the CFA movement subpopulation during the preparation period (Fig. 2c), and the narrow waveforms of the disinhibited neurons upon RFA inhibition (Fig. 7a), the EM results suggest that PV+ interneurons may cancel incoming excitation in the CFA during the preparation period. This assigns the RFA a major role in inhibiting involuntary movements by modifying CFA local circuits.

Fourth, we provided new insights into the mechanisms and circuit dynamics between the RFA and CFA for gating movements. Previously, it has been shown that the information from the RFA is important for adapting CFA activity appropriately to task demands during locomotion on differently spaced ladder rungs^30^. The activity in the RFA also displays high specificity to either internally generated or externally triggered movements^8,47^. In line with our results, this context-dependent information is not conveyed consistently to the CFA but can vary over behavioral sessions, and correlates with behavioral performance, at least in axons targeting superficial layers^47^. With our data set we close the knowledge gap about which kind of information is conveyed by neurons projecting from RFA to CFA during the distinct phases of motor planning and movement execution and how these neurons affect CFA activity during these different epochs. Specifically, we found differential impacts of RFA inputs on the CFA depending on the cortical layer as well as the context (i.e., activity outside the task compared to the task phases preparation and movement). During rest, RFA input to the CFA exclusively affected the deep layers in a predominantly excitatory manner. During the preparation period of the task, neurons projecting from RFA to CFA exhibited sustained activity that predominantly increased the activity of the neurons in the superficial layers and decreased the activity of the neurons in the deep layers of the CFA. During the movement period, neurons projecting from RFA to CFA showed transient activity that was linked to an increase in activity in the superficial layers, potentially via indirect pathways^13^, and bidirectionally modulated neurons in the deep layers of the CFA. Previously, it has been suggested that PV+ interneurons in the CFA play a vital role in the execution of movement^48^ and are activated before pyramidal cells during reaching^44^. Here, we propose that, depending on the task phase and the subpopulations that are active in RFA, RFA inputs engage inhibitory and excitatory neurons in the CFA to different extents to modulate local recurrent circuits. In line with a study combining optogenetics and fMRI^49^, the excitation of superficial layers argues for the upper layers having a role in local motor cortex computations by selectively activating ensembles in deep layers, generating the desired motor output^25^ (Fig. 9). Recurrent circuits may selectively amplify certain patterns in the feedforward input, enhancing the signal-to-noise ratio of the selected patterns^50,51^. Thereby, small patterned fluctuations in the difference between excitation and inhibition will drive large patterned fluctuations in the sum of excitation and inhibition^52^. This might partly explain the context dependent and delayed effects: depending on the state of CFA, incoming RFA inputs are amplified differentially depending on the ongoing local circuit activity and the balance between local excitation and inhibition. Through the lens of dynamical systems^2^, this could be phrased as follows: The pathway specific manipulations cause changes in the CFA population along a behaviorally relevant manifold as the RFA to CFA connection is physiologically linked to motor behavior. Depending on the task phase, the behaviorally relevant manifolds in CFA and their relation to the RFA inputs differ, thus impacting the effect of the optogenetic manipulations.

Our study allows addressing the longstanding challenge of interpreting mixed brain activities within brain areas where subpopulations dedicated to specific tasks are intermingled. The newly developed NeuRL tool enables the deciphering of the neural code of specific subpopulations and is amendable to humans, allowing a direct translation of our results to clinical applications. It solely requires extracellular neuronal measurements—a technique which was established in the 1950s and developed continuously in human patients up to modern high-density probes^53–58^. Thus, our newly gained knowledge about the role of specific neuronal ensembles can be incorporated into the design principles of modern brain-computer interfaces^59–66^ to allow even better control of external devices.

## Acknowledgements

We would like to thank Sigrun Nestel for technical assistance in electron microscopy, Dan O’Shea and Andreas Draguhn for comments on an earlier version of this manuscript, and Zongpeng Sun and Philippe Coulon for discussions about the experimental design and follow-up experiments, respectively. We thank Etienne Ackermann and scidraw.io for the illustrations. This work was supported by the Bernstein Award 2012 (01GQ2301), the BrainLinks-BrainTools Cluster of Excellence (EXC 1086), the Deutsche Forschungsgemeinschaft (DFG) via Grants DI 1908/3-1, DI 1908/11-1 and DI 1908/6-1, and the ERC Starting Grant OptoMotorPath (338041), which were all awarded to I.D.

## Author contributions

I.D. and J.B. conceived the study. M.A., I.D., G.K., J.A, J.B., and A.V. wrote the manuscript. M.A. performed all in vivo experiments, excluding those involving electron microscopy, and performed the data analysis. G.K (Gabriel Kalweit), M.K., and J.B. designed the Q-learning approach. G.K. and M.K. performed the in silico experiments. G.K. (Golan Karvat) designed and built the electrophysiological and behavioral setup. A.S. supported the histological evaluations. A.A. supported the confocal imaging.

## Online Methods

### Animals

Female adult CD® IGS rats (Sprague Dawley; 11 rats; 8 weeks of age; 250–300 g) were used in this study. They were group-housed (2 to 4 per group) in large double-decker cages under a reversed 12 h light-dark cycle (light off from 8 a.m. to 8 p.m., for the duration of the training and experiments). One week before the first behavioral training, rats were handled daily. Food (standard lab chow) and water were provided ad libitum. During the course of the experiment, free access to food was maintained but water was restricted while keeping the rats at > 85% of their weight before water restriction. For 2 days per week, ad libitum food and water access were ensured. All animal procedures (surgeries, behavioral training, optogenetics, electrophysiological recordings, and perfusions) were approved by the Regierungspra □ sidium Freiburg, Germany.

### Behavioral setup

We developed a delayed Go/No-Go task and setup for freely moving rats^10^. In this study we focused on the Go part of the task. Early training utilized 4 custom-built setups for simultaneous training controlled individually by Med-PC software (Med Associates, Fairfax, VT). Each training box included a 30 × 25 × 30 cm Plexiglas box with a grounded metal floor. A 2 × 12 mm infusion cannula (1464LL, Acufirm, Dreieich, Germany) covered with a 7 mm (diameter) metal ball served as a lever. The lever was clamped to a holder 1 cm above the cage ground and between two 65 × 35 mm retractable Plexiglas walls. The holder was centered by two pairs of magnets (adhesive force 2.5 kg, model S-08-08-N, Supermagnete, Gottmadingen, Germany). We controlled the distance between magnets, and thus their force, by attaching them to M12 screws. The axis of the holder was connected to a 10-bit magnetic angle encoder (AEAT-6010, Avago Technologies, San Jose, CA) reporting the left-right position. The metal holder was connected to a 5V source, thus forming a conductive touch sensor. To deliver vibrotactile stimuli, we glued a small vibrating motor (3V ERM motor, Digikey no. 1597-1244-ND, Seeed Technology Co., Shenzhen, China) to the lever. The vibrator, touch sensor, and angle encoder were controlled by an Arduino Uno (Arduino, Turin, Italy) connected via transistor-transistor-logic (TTL) to the Med Associates control cabinet. We controlled the red cage light, reward delivery infusion syringe pump (PHM-107, Med Associates), and cage speaker (ENV-224AM, Med Associates) directly with the Med Associates cabinet.

### Behavioral training

We initially trained the rats to hold the lever steadily. First, we acclimated the rats to the behavioral setup for one 30 min session. Then, we rewarded the rats with 3% sucrose water accompanied by a 12 kHz tone clicker upon touching and/or moving the lever. After associating touching the lever with the reward, we used Plexiglas walls to restrict the rats from using body parts other than the forepaw. In gradual steps, we narrowed the gap between the restriction walls to 2 cm and automatically increased the holding duration in steps of 10 ms after each successful hold. If a rat used its mouth or both forepaws, we manually decreased the reward size, otherwise, we increased the reward size following successful trials. We inspected the preferred holding direction and paw of each rat. A hold was defined as moving and keeping the lever beyond a 1 mm threshold. Typically, the rats pulled the lever in the preferred horizontal direction to ∼5 mm until reaching a mechanical limit.

Next, we introduced the vibrotactile stimulus. To discourage timing, we randomized the holding duration until stimulus presentation between 600 and 1,600 ms with a 1:1 ratio (Fig. 1), and the allowed reaction time (RT) window was automatically decreased from 2,000 to 600 ms. The stimulus frequency was set to ∼200 Hz. Throughout the behavioral task, we used the 12 kHz tone clicker to indicate correct trials to the rats and white noise to indicate errors (early/late releases). After an error, at least 1 s had to pass with the lever at center until a new trial could begin (time-out).

### Stereotaxic injections and implantation sites

The animals were initially anesthetized with isoflurane inhalation, followed by intraperitoneal injection of 75 mg/kg ketamine (Medistar, Holzwickede, Germany) and 50 mg/kg medetomidine (Orion Pharma, Espoo, Finland). The animals were put into a transportation container covered with an opaque cloth to facilitate the anesthetization. Once the animals were anesthetized, they were positioned in a stereotaxic frame (David Kopf Instruments, Tujunga, CA, USA), and their body temperature was maintained at 36–37 °C using a rectal thermometer and a heated blanket (FHC, Bowdoin, USA). The animals were kept anesthetized using ∼1.0% isoflurane and 1.0 l/min O_2_. For pre-surgery analgesia, we subcutaneously administered 0.05 mg/kg buprenorphine (Selectavet, Dr Otto Fischer GmbH, Weyarn/Holzolling, Germany). Every hour, we injected 2 ml of isotonic saline subcutaneously. We applied a moisturizing ointment to the eyes to prevent them from drying out (Bepanthen, Bayer HealthCare, Leverkusen, Germany). Their skin was disinfected with Braunol (B. Braun Melsungen AG, Melsungen, Germany) and Kodan (Schülke, Norderstedt, Germany) using sterile cotton tips. To perform the craniotomy, a 2 cm-long incision of the skin on the head was opened using a scalpel. The exposed bone was cleaned using a 3% peroxide solution. Craniotomies were drilled bilaterally extending from −2 to +4.5 mm in the anterior-posterior direction and +1 to +4 mm in the lateral-medial direction relative to Bregma.

For phototagging experiments (N = 3 rats; table 4), we injected a 1 _μ_l viral vector (rAAV5/hSyn-hChR2(H134R)-eYFP, UNC Vector Core, Chapel Hill, North Carolina) into the RFA at two different depths (0.6 and 1.2 mm, 0.5 _μ_l each). For experiments on the inhibition of neurons projecting from RFA to CFA, we injected 1.4 _μ_l of viral vector (rAAV2-Retro/CAG-Cre, UNC Vector Core, North Carolina, Chapel Hill) at four different sites in both hemispheres of the CFA (two points separated by ∼0.5 mm in the anterior-posterior axes at depths of 0.6 and 1.2 mm, 0.35 _μ_l at each point). We injected two sites in the RFA with 0.75 _μ_l of viral vector in both hemispheres (rAAV5/EF1a-DIO-eNpHR3.0-eYFP, UNC Vector Core) at depths of 0.6 and 1.2 mm (0.35 _μ_l at each point; table 5). We injected the respective areas at a rate of 100 nl/min using a 10 _μ_l gas-tight Hamilton syringe (World Precision Instruments, Sarasota, Florida). To minimize reflux of the injected volume, we left the injection needle in the tissue for 10 additional minutes before slowly extracting it from the brain.

For electrophysiological recordings, we inserted 2-shank, 32-channel laminar probes (E32+R-150-S2-L6-200-NT, ATLAS Neuroengineering, Leuven, Belgium) into the contralateral hemisphere relative to the forepaw used in the experiment (N = 5 rats each with probes in the RFA or CFA, N = 10 rats overall; Extended Data Table 5). The probe was slowly inserted into the brain while the rat was held with a vacuum holder (ATLAS Neuroengineering, Leuven, Belgium).

We applied a sealant (Kwik-Cast, World Precision Instruments) over the craniotomy and fixed the probe and/or optical fibers to the skull with UV-cured dental cement (RelyX, 3M, Saint Paul, MN). Self-tapping skull screws (J.I. Morris Company, Southbridge, Massachusetts) acting as a reference for extracellular recordings were placed above the cerebellum. For increased stability and reduced noise, we also cemented the custom-made electrode interface board. Rats were given >7 days of recovery before the continuation of experiments.

### Data acquisition

We performed electrophysiological recordings during 3–4 sessions (minimum 15 correct trials) per rat with at least one week break in between sessions. We sampled the broadband signal at_∼_25 kHz using a digital head stage (ZD32, Tucker-Davis Technologies (TDT), Alachua, Florida). Spiking activity was bandpass filtered between 300 and 5,000 Hz. The Med Associates system registered behavior at 100 Hz and synchronized with the electrophysiological signal via TTL communication.

Phototagging experiments were conducted eight weeks after the viral injection. Laser stimulation was conducted at 473 nm with a power of 1, 2, 4 or 8 mW out of a 200 µm optical fiber with at least 30 trials for each power setting at the end of the behavioral sessions during continued neuronal recordings (N = 3 rats, 1 session per rat).

For inhibition experiments, we inhibited the RFA during either the preparation or movement periods. We used light with a wavelength of 590 nm at 15–20 mW for 0.5 s or until movement initiation (50% of trials). After the behavioral session, we inhibited for 0.5 s with an interstimulus period of 10 s.

## Data Analysis

### Behavior

RTs were computed from the stimulus onset until the rat released the lever. Lever release was defined as the movement of the lever laterally to an amplitude of at least 2 mm. In the RT distributions plot (Fig. 1d), RTs were grouped in 10 ms bins for visual display. Lever position was quantified during the preparation period in an analysis window from 0.3 s before release up to release onset.

### Electrophysiology

We sorted the broadband signal into units using KiloSort ^67^, inspected each cluster, and defined units based on wave shape. The binary spike-time array (1 for spike, 0 otherwise) of each unit was smoothed into FR with a Gaussian kernel with a standard deviation of 50 ms and then normalized (z score, baseline = 3 to 2 s before trial start). We defined units as modulated if the absolute z value of the FR in correct trials crossed a threshold of 1.96.

To classify neurons, we used a metric that compares the mean firing rate during the preparation and movement periods:

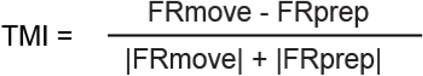

with FRmove referring to the firing rate during the movement period from the go cue to 1 s after the go cue and FRprep referring to the firing rate during the preparation period from trial initiation to the go cue. If TMI > 0, a neuron was classified as a movement neuron; otherwise, it was classified as a preparation neuron.

In phototagging experiments, a neuron was considered as ‘tagged’ if it fulfilled three criteria: (1) a light pulse of 1 ms caused an action potential in the recorded single unit for at least 60% of the laser pulses (Fig. 4d top); (2) the light-induced action potential waveform was similar to the mean of the spontaneously occurring waveforms with a Pearson correlation coefficient higher than 0.7 (Fig. 4d inset), and (3) the induced action potential occurred within a latency window of 15 ms, which was in line with the timing of antidromically-evoked action potentials ^68^ (Extended data Fig. 5).

For inhibition experiments, we evaluated the performance (i.e., the difference in error rate) as well as the effect on RT when inhibiting during the preparation or movement periods. We only considered early releases to be those occurring 0.5 s before the go cue to produce data comparable to those from early trials with a laser, which started 0.5 s before the go cue. We conducted the RT analysis for 10 consecutive sessions per rat.

### Histology

For the histological verification of eNpHR3.0, we euthanized the animals by administering 400 mg/kg sodium pentobarbital (Release®500, WDT, Garbsen, Germany) and transcardial perfusion with PBS followed by ice-cold 4 % paraformaldehyde. After removing the brains, we post-fixed the tissue for 2 additional days before transferring it into a solution of 30 % w/v sucrose (Merck KGaA, Darmstadt, Germany) in water at 4 °C for equilibration. Brains transferred from a sucrose solution were attached to the cooling block of a microtome with Tissue Tek (Sakura Finetek, Fisher Scientific, Germany) and were sectioned into 50 _μ_m thin slices. The slices were transferred to phosphate-buffered saline (PBS) with 0.01% sodium azide. For antibody staining, selected slices were washed for 3 × 10 minutes in PBS on a rotary shaker at room temperature. The slices were blocked and permeabilized for 1 hour (PBS 0.01 M/Triton 0.4%/BSA 5%, Sigma Aldrich, St. Louis, MO, USA) on the rotary shaker. The first antibody (dilution 1:1,000, monoclonal anti-Parvalbumin, P3088, Merck, Taufkirchen, Germany) was applied overnight at 4 °C (PBS 0.01 M/Triton 0.2%). The slices were washed for 3 × 10 min in PBS on the rotary shaker at room temperature. The second antibody (dilution 1:250, Cy3 goat anti-mouse, M30010, Thermofisher, Waltham, MA, USA) was applied for 3 h (PBS 0.01 M/Triton 0.2%). Finally, the slices were washed for 3 × 10 min in PBS on the rotary shaker at room temperature and mounted. The slices were imaged with a Zeiss (Oberkochen, Germany) LSM880 confocal microscope using a 40x objective.

### Electron Microscopy

The protocol for EM imaging has been described previously^69^. Briefly, slices were fixed in 4% PFA (w/v in 0.1 M PB; Polysciences Europe GmbH, Hirschberg a.d. Bergstraße, Germany) and 2.5% glutaraldehyde (w/v in 0.1 PB; Carl Roth GmbH, Karlsruhe, Germany) overnight. After fixation, slices were washed for 4 hours in 0.1 M PB. Subsequently, slices were incubated with 1% osmium tetroxide (Carl Roth) for 45 minutes, washed in graded ethanol (up to 50% [v/v]) for 5 minutes each, and incubated with uranyl acetate (1% [w/v] in 70% [v/v] ethanol; Science Services, Munich, Germany) overnight. Each slice was then dehydrated in graded ethanol individually (80%, 90%, 98% for 5 minutes, 2 100% for 10 minutes). Subsequently, the slices were washed in propylene oxide (Polysciences Europe GmbH) twice for 10 minutes before incubation with durcupan/propylene oxide (1:1 for 1 hr; Sigma-Aldrich, Taufkirchen, Germany) and transferred to durcupan (overnight at room temperature). Slices were embedded in durcupan and cut into ultra-thin sections (55 nm) using a Leica UC6 Ultracut (Wetzlar, Germany). Sections were mounted onto copper grids (Plano, Wetzlar, Germany), and an additional Pb-citrate (Carl Roth, Karlsruhe, Germany) contrasting step was performed (3 minutes). Electron microscopy was performed using a Philips CM100 microscope equipped with a Gatan Orius SC600 camera (Gatan, Pleasanton, CA, USA) at 3,900 magnification. Acquired images were saved as TIF files and analyzed by an investigator blinded to experimental conditions.

### Q-learning and hypothesis generation Inverse reinforcement learning

We modeled the task of neuronal decoding in the reinforcement learning framework as a Markov Decision Process (MDP), where an agent (a rat) acted in an environment (preparation-movement task). Following policy π by applying action *a*_*t*_ ∼ π from *n*-dimensional action-space *A* in state *S*_*t*_, the subject reaches a state *S*_*t*+1_ ∼ *M* according to the stochastic transition model *M* and receives scalar reward *r*_*t*_ in each discrete time step *t*. The agent has to adjust its policy π to maximize the expectation of long-term return *R* (*S*_*t*_) = Σ_*t*’ ≥*t*_ γ^*t*’−*t*^ *r*_*t’*_, where γ ∈ [0,1] is a discount factor which can be used to give more recent rewards a higher weight and to prevent the long-term value from running infinite in non-terminal problems (in our experiments, we use γ =1, i.e. no discount). The action-value function represents the expected long-term value of an action when following policy *π*(i,e., 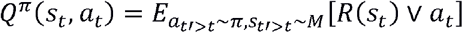. From the optimal action-value function *Q*^*^ one can derive a corresponding optimal policy *π*^*^by maximization. Inverse reinforcement learning recovers a reward function from observed trajectories from expert policy π^*R*^ under the assumption that the agent was softly maximizing the induced expected long-term return (i.e., according to a probability distribution). This problem has been solved previously using different approaches such as Max Entropy IRL^70^, which could be costly and lead to approximation errors when estimating the scalar immediate reward.

### Action-value Iteration

Here, we focus on finding an optimal policy via model-based action-value iteration. The Q-function, represented by a table with entries for every state and action, gets updated in every iteration *k* based on the Bellman optimality equation with a given transition model *M*:

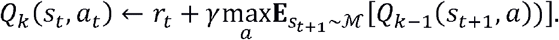

### Formalization

In this section, we describe how to infer the scalar underlying reward function of a rat’s behavior, the supervised approximation of this scalar reward as a weighted combination of neuronal signals, and the neuronal decoding mechanism using the intrinsic reward function.

### Estimation of Intrinsic Reward

Our main assumption is that the rodent is softly (i.e., according to a probability distribution) maximizing its measure of optimality, which we define to be the expected cumulative sum of an unknown immediate reward function, also known as the Q-value. The Q-value likely corresponds to activity in brain regions responsible for planning and movement^71^. The assumption of soft maximization of the measure of optimality is known as the Boltzmann assumption and it has already been applied to model the behavior of humans and animals in a plethora of prior studies^72–74^. In other words, the actions taken by the animal are samples from a Boltzmann distribution over its optimal action-values *Q* ^*\**^(*S*, ·):

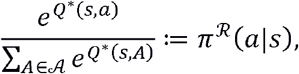

We assume the rodent to softly maximize its measure of optimality which we define to be the expected cumulative sum of an unknown immediate reward function; that is, the actions taken by the rat were sampled from a Boltzmann distribution over its optimal action-values *Q* ^*\**^(*S*, ·):

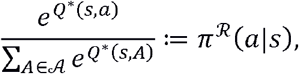

for all actions *a* ∈ *A*, and concomitantly:

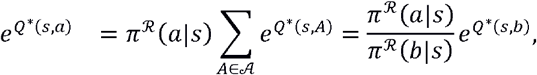

for all actions *b* ∈*A*_*á*_ where A_*á*_ = A{*a*}. Following the derivations as proposed previously ^75^:

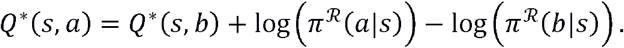

Using the Bellman optimality equation, the immediate reward of action a in state s could thus be expressed by the immediate reward of some other action *b* ∈ *A*_*á*_. The respective log-probabilities and future action-values are:

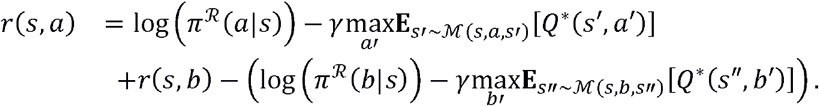

Substituting the difference between the log-probability and the discounted action-value of the future state *s’* as:

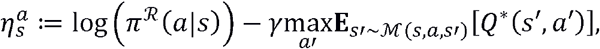

We could put the reward of action *a* in state *s* in relation to the reward of all other actions:

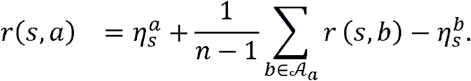

The resulting system of linear equations could be solved with least squares. We started by estimating the immediate reward for all terminal states and then went through the MDP in reverse topological order based on the model *M*. The Boltzmann distribution induced by the optimal action-value function on this learned reward was equivalent to the demonstrated arbitrary behavior distribution ^75^. Inverse action-value iteration (IAVI) thus returned a scalar intrinsic reward function which precisely encoded the recorded behavior of the rats as an intermediate result that served as a supervised signal to learn new features from neuronal spiking.

Note that while we exploit the stochastic behavior assumption based on the Bellman optimality equation and while there is a connection between the response of dopamine neurons and temporal-difference learning ^76^, we only leveraged the defined computational model to estimate the expected long-term reward without assuming a similar mechanism in the rodent’s brain.

### Mapping of Neuronal Spiking to Intrinsic Reward

As a second step, we mapped the neurons projecting from RFA to CFA to intrinsic reward function in a regression step to draw conclusions about the behavior based on neuronal activity. We hence assumed the immediate reward function to be a projection:

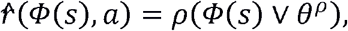

where *ρ* is a parameterized function of features with parameters *θ*^*ρ*^ (e.g., a linear combination or a neural network). The vector *Φ* (*s*) = (*Φ*_1_ (*s*),…, *Φ* _*m*_(s))^T^ contained m features based on the recordings of m neurons, such as the mean activity over all trials. We can fit parameters *θ*^*ρ*^ according to the class of function approximator (e.g., either by least squares or gradient descent) on the difference between reward *r*(*s, a*) and prediction 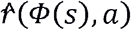. The mapping can then be used to predict the resulting behavior based on neuronal spiking in different trial outcomes.

### Neuronal Decoding from Intrinsic Reward

The parameters *θ*^*ρ*^ of 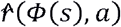 were fitted to represent immediate reward *r* (*S,a*)and hence the underlying behavior of the rats as closely as possible. The resulting parameters could contribute to the generalization to any arbitrary neuronal spiking *Ψ* (*S*) = (*Ψ*_1_(*S*), …, *Ψ*_*m*_ (*S*))^T^ which yields adjusted reward and action-values in each time step *t*:

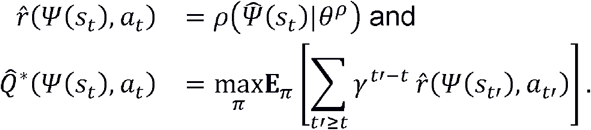

From the optimal Q-function *Q*^*^ (Ψ (*S*),*a*) based on features, *Ψ* (*s*), we infer the respective predicted action-probabilities by:

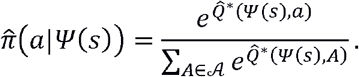

To identify neurons with particular relevance for a specific type of response, we can modulate their neuronal activity by modifying the respective features *Φ*_*i*_(*S*), |_1 ≤*i≤m*_and keeping all other features fixed. Thus, we can simulate the inhibition of certain neurons within the model and make predictions about the trial outcome and RTs. The change in behavior between the ground truth based on the recorded spiking and the predicted behavior based on the modulated features provided insight into the possible individual impact of these neurons on behavior.

### MDP Formulation

We modelled the preparation-movement task as a Markovian decision process (MDP). The MDP was defined as a four-tuple ⟨*S,A,M,r*⟩ where the set of states was defined by *S*={0.0s,0.2s, …,1.2s} ∪ {Before Cue, Cue, After Cue, After Cue1, After Cue2, …, Time to Release, Late Release} ∪{Success,Failure}, discretizing the time into chunks of 0.2 s. In every state, the rat could pick an action from the action space *A* = {stay,release} (Fig. 5). We defined the MDP to have deterministic transitions. In these experiments, we considered the reward function *r* :*S* ×*A* →*R* to be unknown.

We used NeuRL to make predictions about the rats’ behavior when inhibiting different sets of neurons: 1) the neurons projecting from RFA to CFA, including both preparation and movement subpopulations; 2) the preparation subpopulation projecting from RFA to CFA, and 3) the movement subpopulation projecting from RFA to CFA. For each set, we ran 3,000 simulations for control trials (no inhibition; 1,000 trials), inhibition during the preparation period (1,000 trials), and inhibition during the movement period (1,000 trials). We determined whether the simulated trial was early, correct, or late according to the discretized simulated release times. If the trial was correct or late, we used the release time to compute simulated RTs.

### Statistical tests

Data are represented as mean +/-SEM. All statistical analyses were computed in MATLAB (Mathworks, version R2018b). In NeuRL simulations, we used a two-sample t-test and applied the post hoc Holm–Bonferroni method for multiple comparisons by adjusting the P-value correspondingly. The respective exact P-value is given in the Results section. For the comparison of distributions in error trial analysis, we used a two-sample Kolmogorov-Smirnov test. For optogenetics experiments and comparing firing rates across conditions, we used a repeated measure two-way ANOVA. A significant difference between two data sets was assumed when the Holm–Bonferroni-corrected P-value <0.05 (indicated by one asterisk in the figures).

**Extended Data Fig. 1:**
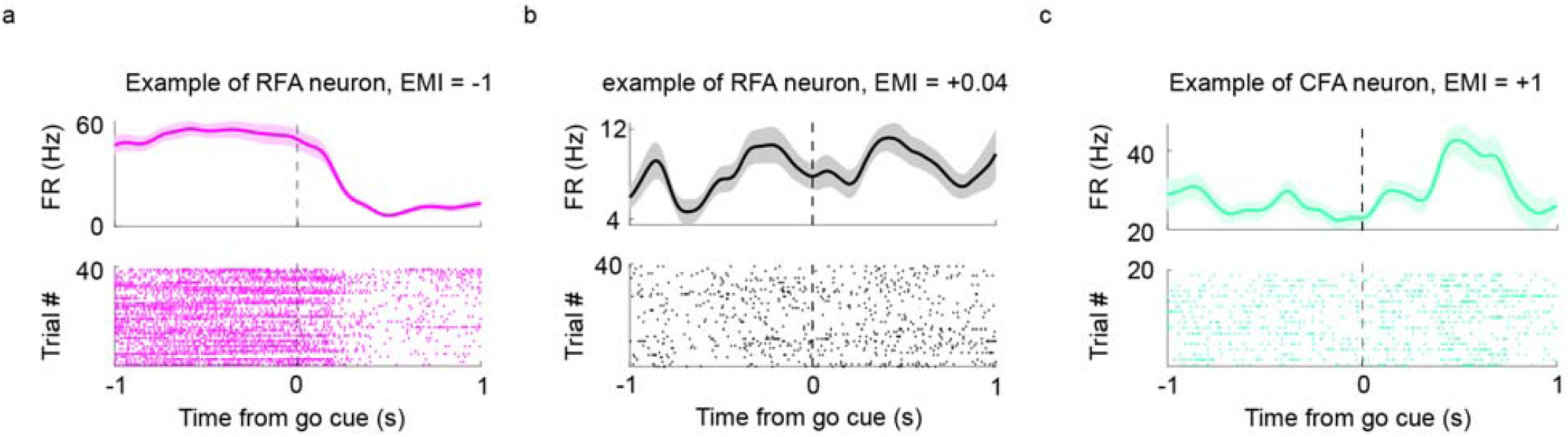
Raster and PSTH of neuron examples with extreme Task Modulation Indices (TMI). Related to Fig.2 **a**. Example of an RFA preparation neuron with a TMI = -1. **b**. Example of an RFA neuron with a TMI close to zero. **c**. Example of a CFA movement neuron with a TMI = +1. The shaded background shows SEM.

**Extended Data Fig. 2:**
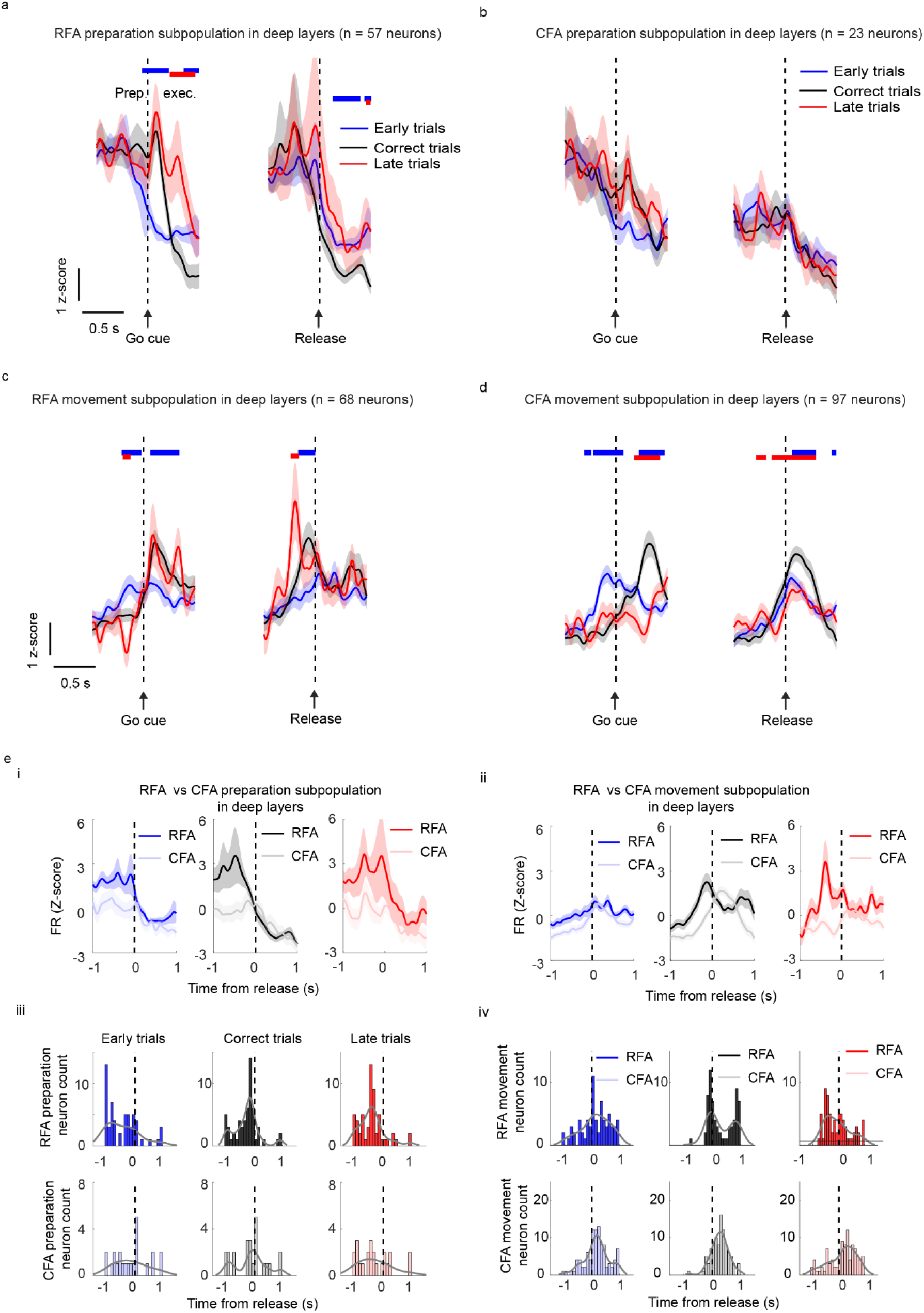
Similar FR patterns in deep-layer subpopulations compared to an analysis of pooled superficial and deep subpopulations (Fig. 3). Related to Fig. 3. The shaded background shows SEM.

**Extended Data Fig. 3:**
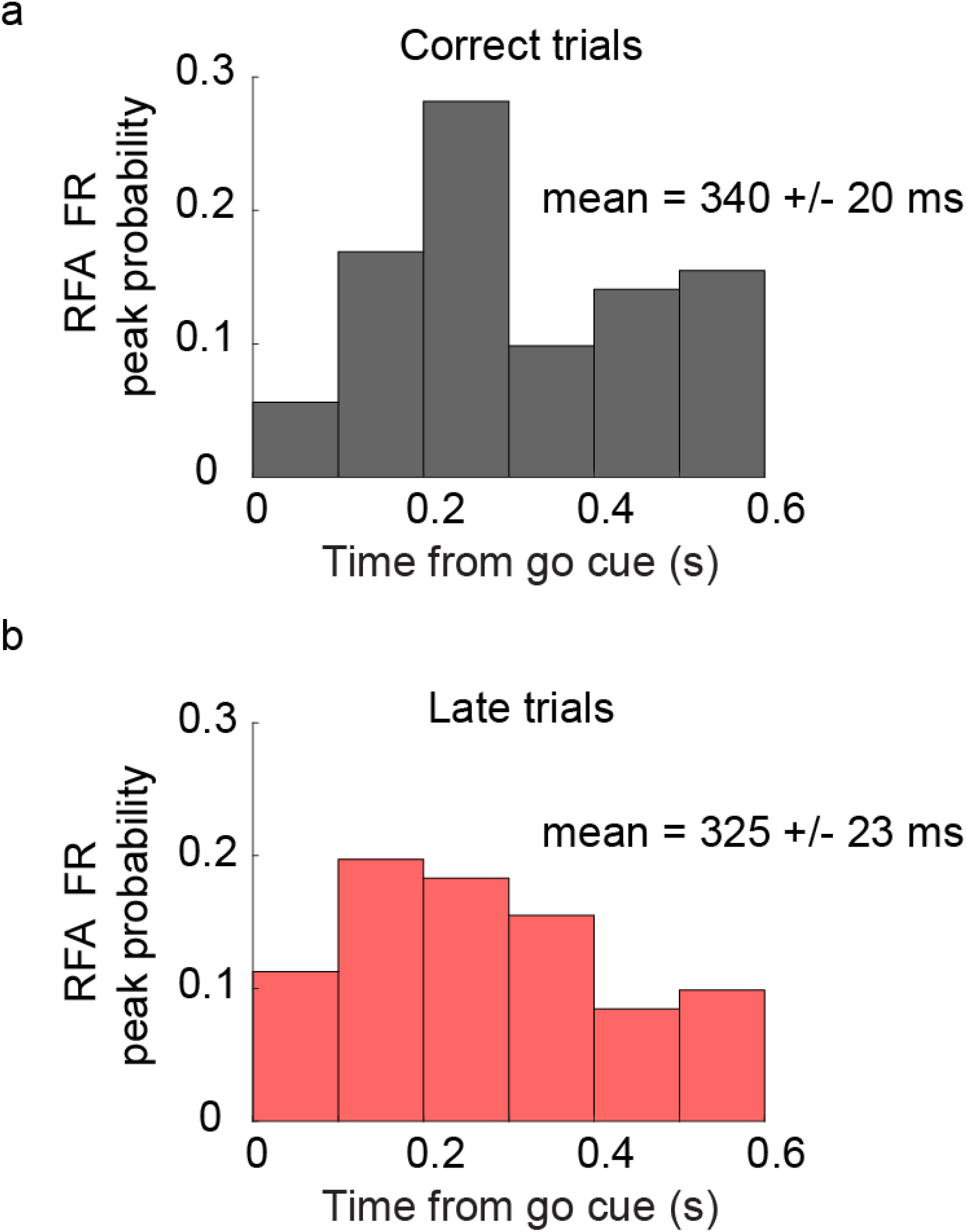
Similar Peak FR latency in RFA in correct and late trials. Related to Fig. 3. **a**. Probability of the peak firing rate of RFA neurons in correct trials. **b**. Same as **a** but in late trials.

**Extended Data Fig. 4:**
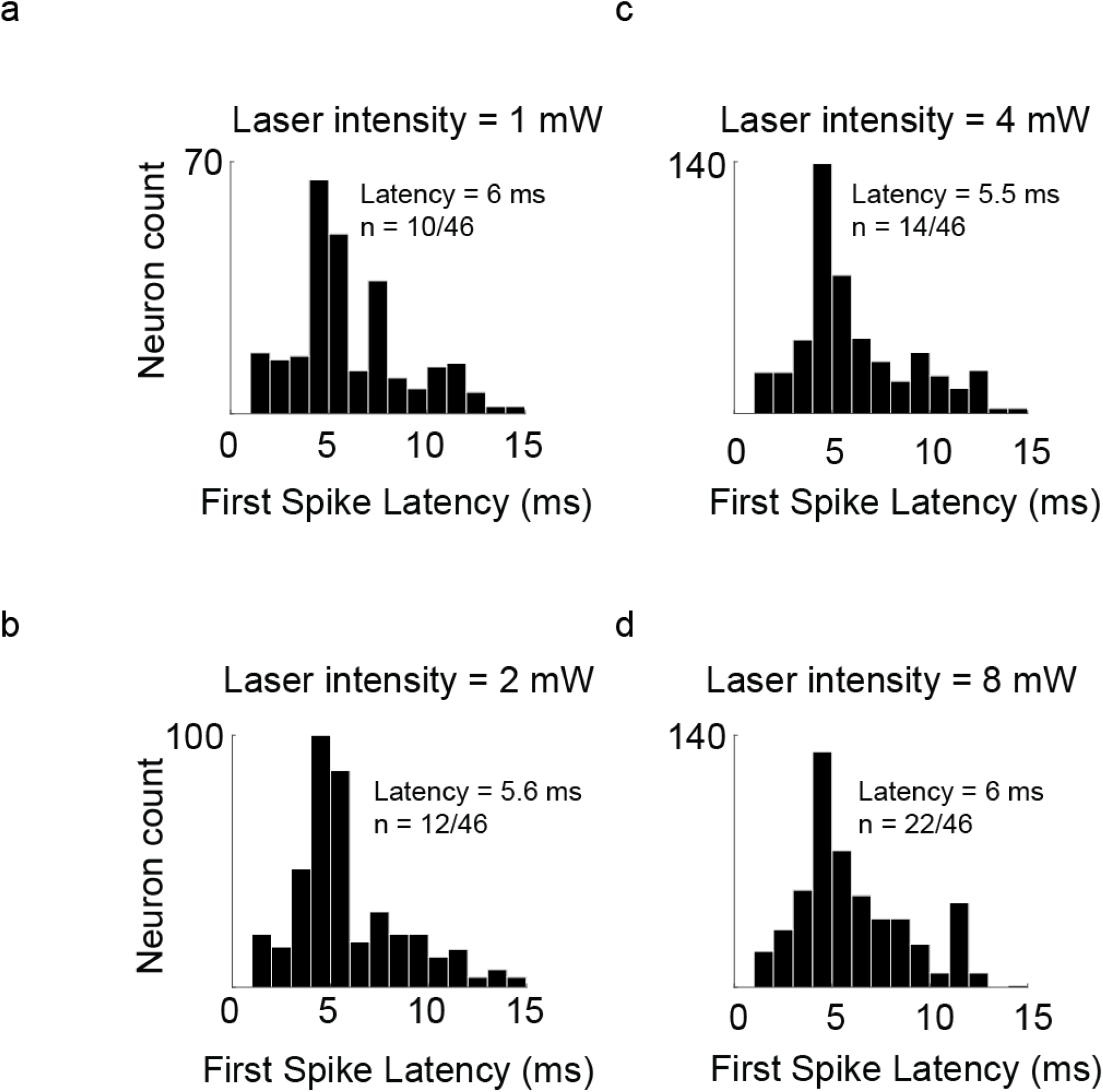
First spike latency of tagged neurons is not affected by laser intensity. Related to Fig. 4. **a**. Population latency response with a laser intensity of 1 mW. **b, c**, and **d** same as **a** but for intensities at 2, r, and 8 mW respectively.

**Extended Data Fig. 5:**
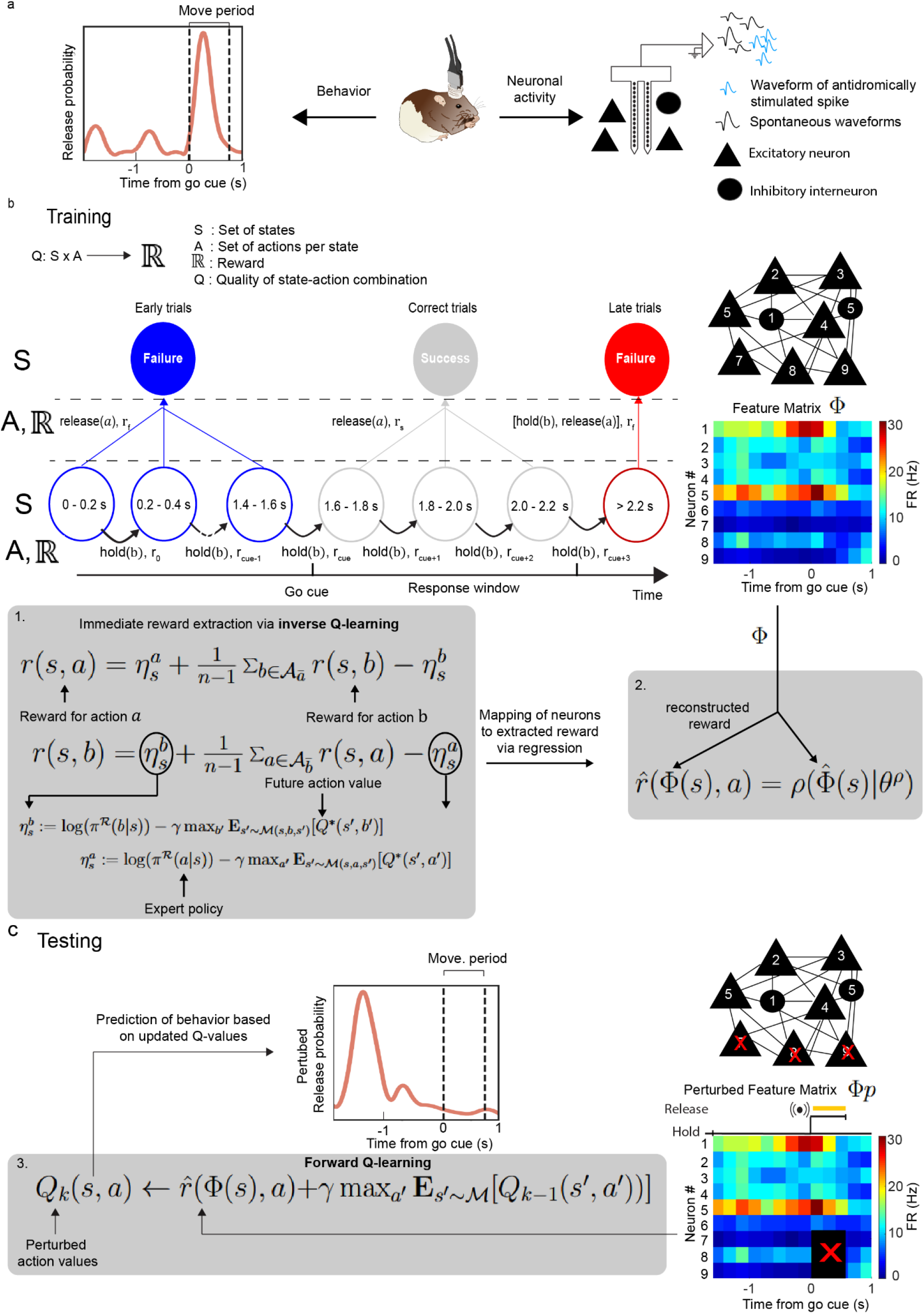
Equations and their relation to the Q-learning flow. Related to Fig. 5 **a**. The inverse Q-learning framework used the rats’ behavior (lever release times) and neuronal activity, consisting of the mean firing rates of RFA neurons in single trials, to assign functions to different neuronal subpopulations (N = 3 rats). Recordings from neurons identified as projecting from RFA to CFA were used to predict the trial outcomes and RTs. **b**. Training the network: a reward value was estimated from actions (hold referring to action b and release referring to action a) and states (success or failure) over 0.2 s bins for each trial. During the preparation period (0 to 1.6 s; blue circles), a state was termed ‘success’ if the action was hold and ‘failure’ when the action was release. In the movement period, a state was termed ‘success’ if the action was release, and after the movement period, a state was termed ‘failure’ regardless of the action. Based on the actions and states, a reward value was extracted via inverse Q-learning. An inverse Action-value iteration (IAVI) is used to compute the immediate reward of action **a** in state **S** can be expressed by the immediate reward of some other action **b** (1), returning an intrinsic reward function that encoded the behavior of the rats as an intermediate result that serve as a supervised signal to learn a mapping from the neuronal signals. Here we used these reward values to map onto a feature matrix reconstructed from the mean firing rate for each RFA neuron in 200 ms bins, giving each neuron a weight that contributed to the computed reward (2). **c**. Testing the network: based on the extracted rewards, different weights were assigned to the neurons in the behavioral task. Perturbation of different neurons in the feature matrix produced a new perturbed reward value that was computed via forward Q-learning (3) and then used to predict the perturbed behavior.

**Extended Data Fig. 6:**
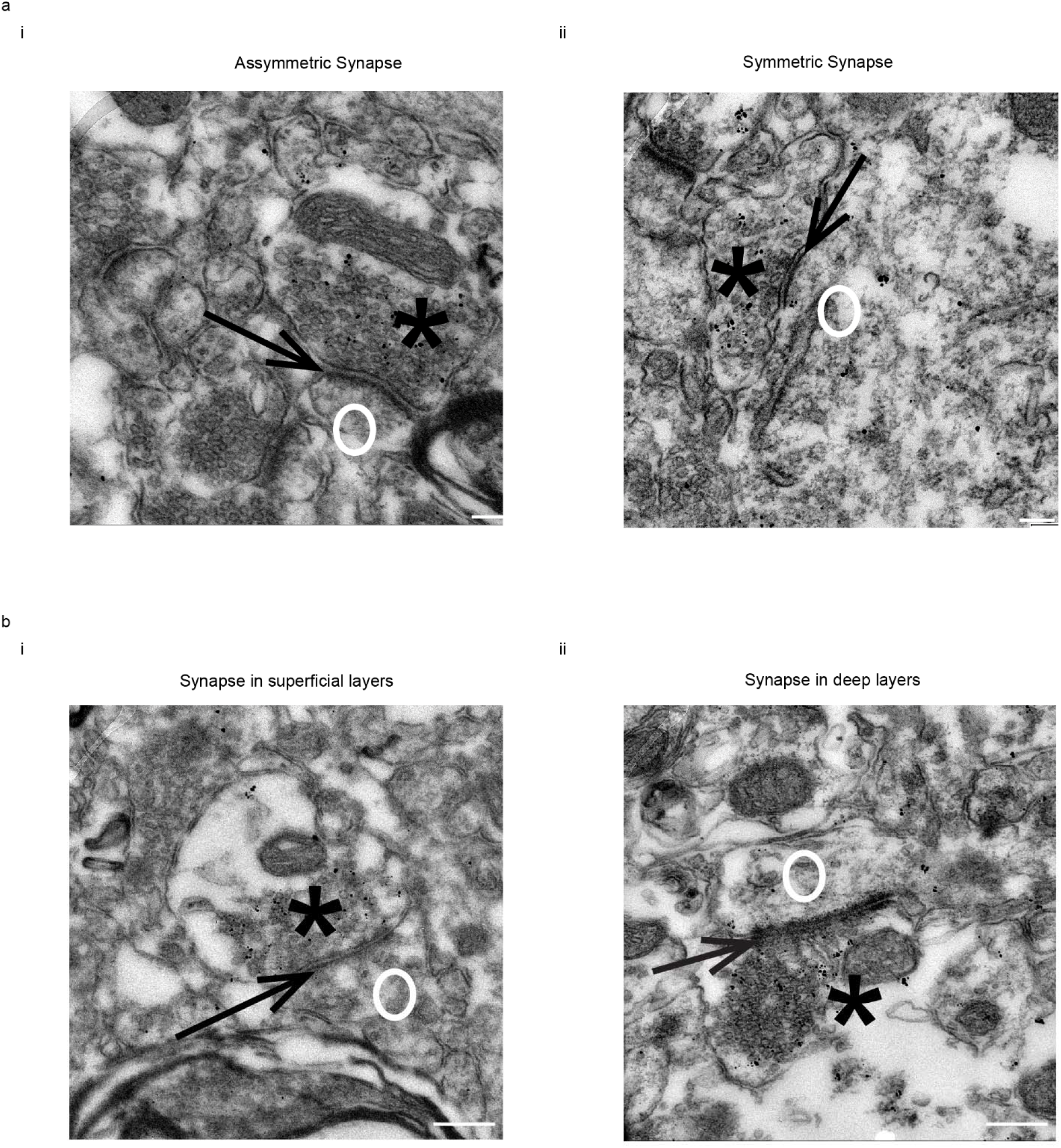
The RFA predominantly forms asymmetric synapses in both superficial and deep layers of the CFA. Related to Fig. 7. **a**. Examples of asymmetric (i) and symmetric synapses (ii). The vast majority of RFA axons form asymmetric synapses in the CFA (97/100, 97%), suggesting that the input from the RFA was dominantly excitatory. **b**. RFA axons target CFA spines in both the superficial (i) and (ii) deep layers. Scale bar in a = 100 nm, scale bar in b = 200 nm, black asterisk: pre-synapse, black arrow: synaptic cleft, white circle: post-synapse.

**Extended Data Fig. 7:**
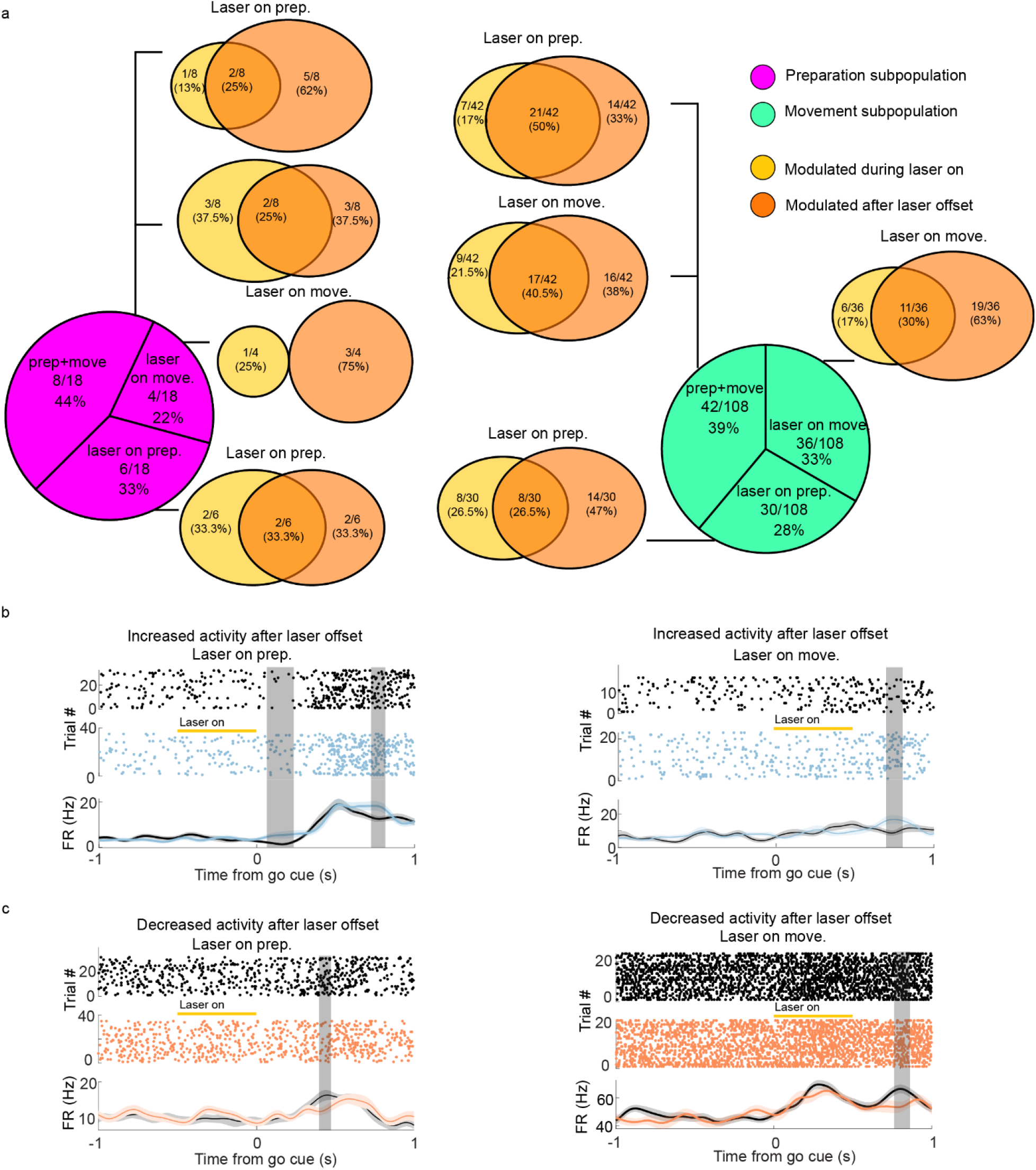
Delayed optogenetic effect on CFA neurons. Related to Fig. 8. **a**. Proportions of preparation neurons (left) and movement neurons (right) modulated by inhibiting neurons projecting from RFA to CFA in different task periods along with fractions of neurons modulated during laser on time, after laser offset, or both. **b**. Raster and PSTH of example neurons with significantly increased activity after the laser offset. Laser was on during preparation period (left) or movement period (right). **c**. Same as b but for neurons with significantly decreased activity after laser offset.

**Extended Data Table 1:**
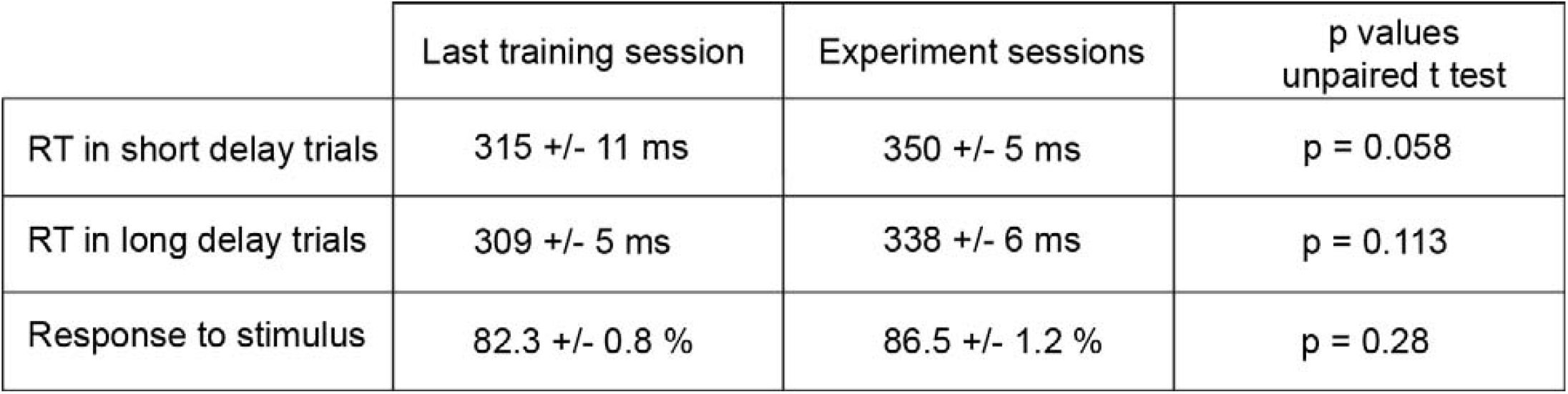
RTs and performance were similar in short- and long-delay trials.

**Extended Data Table 2:** Coordinates of silicon probe implantation sites in the RFA and CFA.

| Rat | Recording area | AP (mm) | ML (mm) | DV of shanks<br>tip (mm) |
| --- | --- | --- | --- | --- |
| 10 | RFA | +3.8 | -1.7 | 2.4 |
| 12 | RFA | +3.7 | -1.5 | 2.4 |
| 504 | RFA | +3.2 | -1.9 | 2.4 |
| 533 | RFA | +3.5 | -2.0 | 2.4 |
| 546 | RFA | +3.5 | -2.0 | 2.4 |
| 1 | CFA | +0.2 | -2.2 | 2.4 |
| 3 | CFA | +0.6 | -2.5 | 2.4 |
| 13 | CFA | -0.1 | -2.1 | 2.4 |
| 16 | CFA | +0.3 | -2.8 | 2.4 |
| 506 | CFA | -0.5 | -2.4 | 2.4 |

**Extended Data Table 3:** Similar peak firing rates in superficial and deep subpopulations.

|  | Superficial subpopulation<br>peak FR (s) | Deep subpopulation<br>peak FR (s) | p value<br>unpaired t test |
| --- | --- | --- | --- |
| RFA early preparation | -0.24 +/- 0.02 | -0.43 +/- 0.07 | 0.63 |
| CFA early preparation | -0.75 +/- 0.28 | -0.14 +/- 0.12 | 0.06 |
| RFA correct preparation | -0.64 +/- 0.2 | -0.27 +/- 0.06 | 0.24 |
| CFA correct preparation | 0.43 +/- 0.1 | -0.12 +/- 0.1 | 0.07 |
| RFA late preparation | -0.01 +/- 0.67 | -0.41 +/- 0.06 | 0.33 |
| CFA late preparation | -0.01 +/- 0.67 | -0.03 +/- 0.12 | 0.49 |
| RFA early move | 0.33 +/- 0.21 | 0.14 +/- 0.06 | 0.52 |
| CFA early move | 0.17 +/- 0.06 | 0.18 +/- 0.04 | 0.88 |
| RFA correct move | 0.01 +/- 0.31 | 0.31 +/- 0.05 | 0.26 |
| CFA correct move | 0.36 +/- 0.04 | 0.27 +/- 0.03 | 0.10 |
| RFA late move | -0.12 +/- 0.22 | -0.04 +/- 0.06 | 0.78 |
| CFA late move | 0.18 +/- 0.10 | 0.16 +/- 0.05 | 0.87 |

**Extended Data Table 4:**
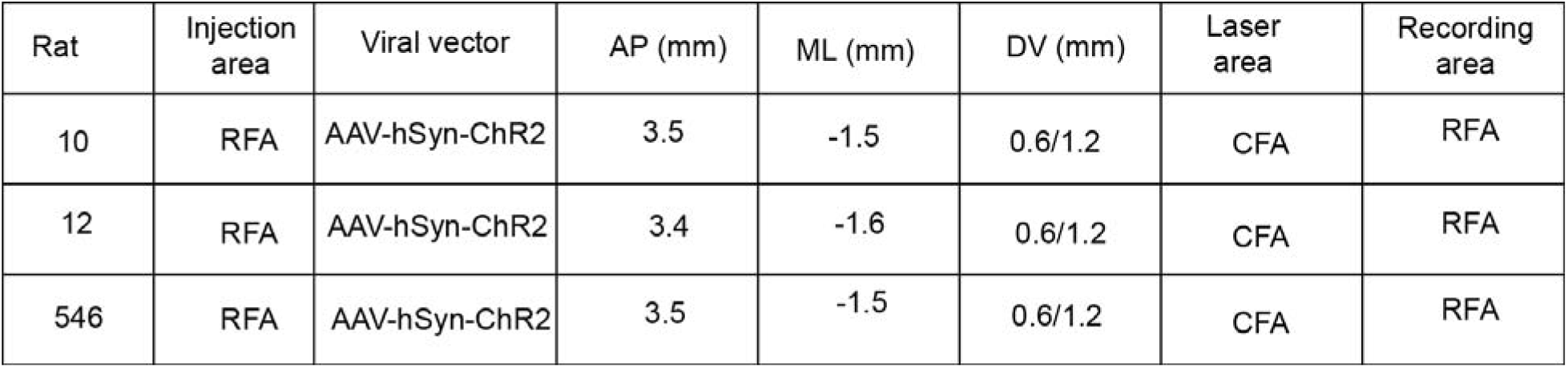
Injection coordinates in phototagging experiments.

**Extended Data Table 5:** Injection coordinates, silicon probes, and optical fiber implantation areas for inhibition experiments.

| Rat | Injection area | Viral vector | AP left (mm) | AP right (mm) | ML left (mm) | ML right (mm) | DV left (mm) | DV right (mm) | Laser area | Recording area |
| --- | --- | --- | --- | --- | --- | --- | --- | --- | --- | --- |
| 3 | RFA | AAV-DIO-eNpHr | 3.7 | 3.6 | 1.5 | 1.7 | 0.6/1.2 | 0.6/1.2 | RFA | CFA |
|  | CFA | retroAAV-cre | 0.5/0.0 | 0.7/0.1 | 2.2/2.3 | 2.5/2.5 | 0.6/1.2 | 0.6/1.2 |  |  |
| 13 | RFA | AAV-DIO-eNpHr | 3.8 | 3.5 | 1.6 | 1.5 | 0.6/1.2 | 0.6/1.2 | RFA | CFA |
|  | CFA | retroAAV-cre | 0.37/0.3 | 0.5/0.0 | 2.8/2.7 | 2.4/2.3 | 0.6/1.2 | 0.6/1.2 |  |  |
| 15 | RFA | AAV-DIO-eNpHr | 3.3 | 3.4 | 1.2 | 1.4 | 0.6/1.2 | 0.6/1.2 | RFA | N/A |
|  | CFA | retroAAV-cre | 0.2/-0.4 | 0.7/-0.1 | 2.6/2.4 | 2.8/2.6 | 0.6/1.2 | 0.6/1.2 |  |  |
| 16 | RFA | AAV-DIO-eNpHr | 3.4 | 3.6 | 1.7 | 1.6 | 0.6/1.2 | 0.6/1.2 | RFA | CFA |
|  | CFA | retroAAV-cre | 0.4/-0.2 | 1.0/0.6 | 2.2/2.3 | 2.6/2.5 | 0.6/1.2 | 0.6/1.2 |  |  |

## Notes

### Competing Interest Statement

The authors have declared no competing interest.

